# Aminoglycoside antibiotics inhibit phage infection by blocking an early step of the phage infection cycle

**DOI:** 10.1101/2021.05.02.442312

**Authors:** Larissa Kever, Aël Hardy, Tom Luthe, Max Hünnefeld, Cornelia Gätgens, Lars Milke, Johanna Wiechert, Johannes Wittmann, Cristina Moraru, Jan Marienhagen, Julia Frunzke

## Abstract

In response to viral predation, bacteria have evolved a wide range of defense mechanisms, which rely mostly on proteins acting at the cellular level. Here, we show that aminoglycosides, a well-known class of antibiotics produced by *Streptomyces*, are potent inhibitors of phage infection in widely divergent bacterial hosts. We demonstrate that aminoglycosides block an early step of the viral life cycle, prior to genome replication. Phage inhibition was also achieved using supernatants from natural aminoglycoside producers, hinting at a broad physiological significance of the antiviral properties of aminoglycosides. Strikingly, we show that acetylation of the aminoglycoside antibiotic apramycin abolishes its antibacterial effect, but retains its antiviral properties. Altogether, this study expands the knowledge of potential aminoglycoside functions in bacterial communities suggesting that aminoglycosides are not only used by their producers as toxic molecules against their bacterial competitors, but could also provide protection against the threat of phage predation at the community level.

## Introduction

Phages are viruses predating on bacteria. Facing the abundance and diversity of phages, prokaryotes have developed multiple lines of defense that can collectively be referred to as the prokaryotic ‘immune system’^1^. Unsurprisingly, phages have evolved a myriad of ways to overcome these barriers, thereby fostering the diversification of bacterial antiviral strategies. Recent bioinformatics-guided screenings revealed a large number of previously unknown antiviral defense systems^2,3^. However, the majority of currently known prokaryotic defense systems relies on a wide range of molecular mechanisms, but is mainly mediated by protein or RNA complexes^4^.

Environmental bacteria produce a wide range of small molecules, conferring producer cells a specific fitness advantage in competitive or predatory interactions. Yet, the potential antiphage role of this extensive chemical repertoire remains largely unexplored. Recently, new types of defense systems have been discovered that rely on small molecules rather than on proteins or RNA^5,6^. Anthracyclines are secondary metabolites naturally produced by *Streptomyces* species and were shown to inhibit infection by double-stranded DNA (dsDNA) phages^5^. These molecules act as DNA-intercalating agents and block the replication of phage - but not bacterial - DNA. Since these secondary metabolites are excreted by *Streptomyces* cells and are diffusible molecules, their production may provide a broad protection against dsDNA phages at the community level.

In nature, producers of secondary metabolites are generally resistant to the molecules they synthetize^7,8^. This feature is of special importance when screening small molecules for antiviral properties, as toxic effects on bacterial growth would prevent the appreciation of any inhibition of phage infection. In this study, we leveraged this principle to look for phage inhibition by secondary metabolites, using bacterial hosts resistant to the compounds tested.

Aminoglycosides are antibiotics well known for their bactericidal effect by targeting the 30S subunit of the ribosome and thereby inhibiting protein synthesis. The aminoglycoside streptomycin, discovered in 1943, was the first antibiotic active against *Mycobacterium tuberculosis^9^*. Decades-old reports described the inhibition of various phages by streptomycin^10–12^. However, the biological significance of these observations was not explored, and the underlying mechanism of action remains unclear. For these reasons, we focused our efforts on aminoglycosides and set to investigate their potential antiphage properties.

In this study, we show that aminoglycoside antibiotics inhibit phages infecting the actinobacterial model species *Streptomyces venezuelae* and *Corynebacterium glutamicum* as well as the λ phage infecting *E. coli*. Investigations of the mechanism of action point towards a blockage of phage infection occurring after DNA injection but before genome replication. Furthermore, the antiphage activity observed with purified aminoglycoside apramycin could be reproduced with supernatants from the natural producer *Streptomyces tenebrarius*, suggesting a broad physiological significance of the antiphage properties of aminoglycosides.

## Results

### Aminoglycosides inhibit a broad range of phages

To investigate a potential antiviral activity of aminoglycosides, we first constructed resistant strains carrying a plasmid-borne resistance cassette encoding an aminoglycoside-modifying enzyme (Tables S1, S2 and S4). We challenged these aminoglycoside-resistant strains with a set of different phages using double-agar overlays with increasing aminoglycoside concentrations as screening platform (Figure 1A). In the screening, we included phages from three different viral realms^13^: dsDNA viruses from the *Caudovirales* order in *Duplodnaviria* (families *Sipho-, Myo*- and *Podoviridae)*, ssDNA viruses from the *Inoviridae* family in *Monodnaviria*, and ssRNA viruses from the *Leviviridae* family in *Ribodnaviria* (Table S3). The efficiency of plating comparing plaque formation under aminoglycoside pressure with aminoglycoside-free conditions was calculated for phages infecting either the actinobacterial model species *Streptomyces venezuelae*, *Streptomyces coelicolor* and *Corynebacterium glutamicum*, or the Gram-negative species *Escherichia coli* (Figure 1B).

**Figure 1.**
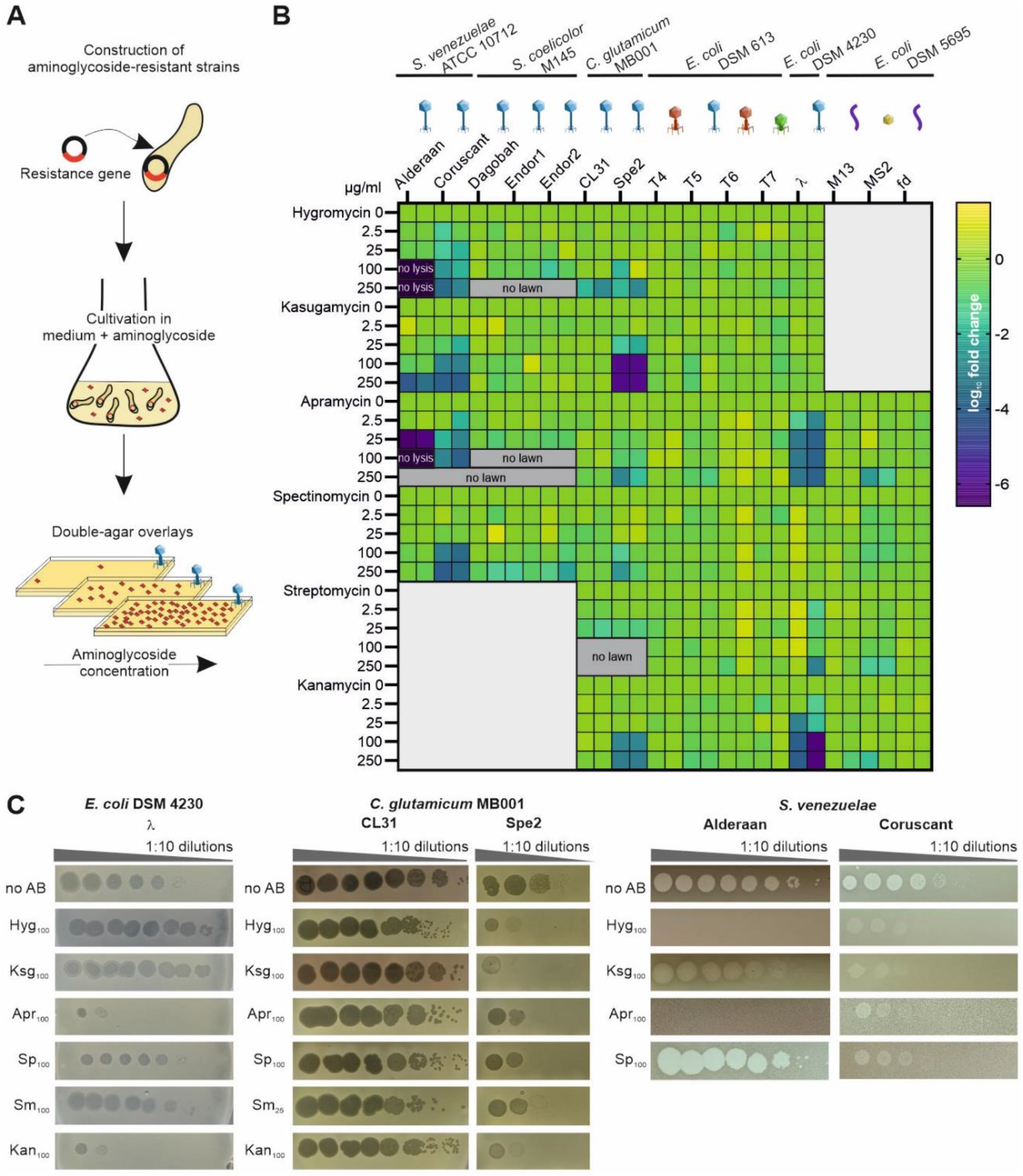
Aminoglycosides inhibit a wide range of phages. **A**, Schematic representation of the screening for the antiphage effect of different aminoglycosides. Strains resistant to the aminoglycosides tested were constructed using plasmid-borne resistance cassettes and subsequently challenged by phages in the presence of increasing aminoglycoside concentrations. **B**, Overview of the screening results, showing the log_10_-fold change in plaque formation by tested phages relative to the aminoglycoside-free control. High concentrations of aminoglycosides prevented in some cases either the formation of plaque or lysis zone by the spotted phages (‘no lysis’) or bacterial growth (‘no lawn’). n=2 independent biological replicates. **C**, Exemplary pictures of propagation assays performed in the presence of the indicated aminoglycoside concentration. Results are representative of two biological replicates.

The extent of inhibition showed clear differences between the individual phages and aminoglycosides. Remarkably, infection with some phages, namely the virulent phages Alderaan, Coruscant and Spe2 as well as the temperate *E. coli* phage λ, was significantly impaired with increasing aminoglycoside concentrations. In contrast, all phages infecting *S. coelicolor,* CL31 infecting *C. glutamicum* MB001 as well as the T-phages, RNA phage MS2 and filamentous phages M13 and fd infecting *E. coli* displayed no susceptibility towards the tested aminoglycosides. The phages susceptible to aminoglycosides infect widely divergent hosts and possess different lifestyles and types of genome ends (Table S3). However, they are all dsDNA phages belonging to the *Siphoviridae* family, suggesting a specificity of aminoglycosides for this phage family.

In the case of *S. venezuelae* phages, we observed the strongest inhibition with the aminocyclitol antibiotic apramycin. The *S. venezuelae* phage Alderaan showed the highest susceptibility among all tested phages, leading to ~10^6^-fold reduction in plaque-forming units for 25 μg/ml apramycin and a complete inhibition of cell lysis at 100 μg/ml hygromycin or apramycin (Figure 1B and 1C). This observation was in line with results from infection assays in liquid culture revealing no more culture collapse when supplementing the respective aminoglycosides (Figure 2A). The antiviral activity was further demonstrated to be dose-dependent, showing already a significant inhibition of infection at 1 μg/ml apramycin (Supplementary Figure S1A). In contrast, only an intermediate inhibition of infection or even no antiviral activity was detected for kasugamycin and spectinomycin, respectively.

To visualize the effect of apramycin on infection dynamics using live cell imaging, *S. venezuelae* mycelium was grown from spores in a microfluidic device and infected with phage Alderaan. Addition of apramycin almost completely inhibited phage-mediated lysis of *Streptomyces* mycelium, confirming the protective effect of apramycin against phage infection (Figure 2B and Supplementary Video S1).

**Figure 2.**
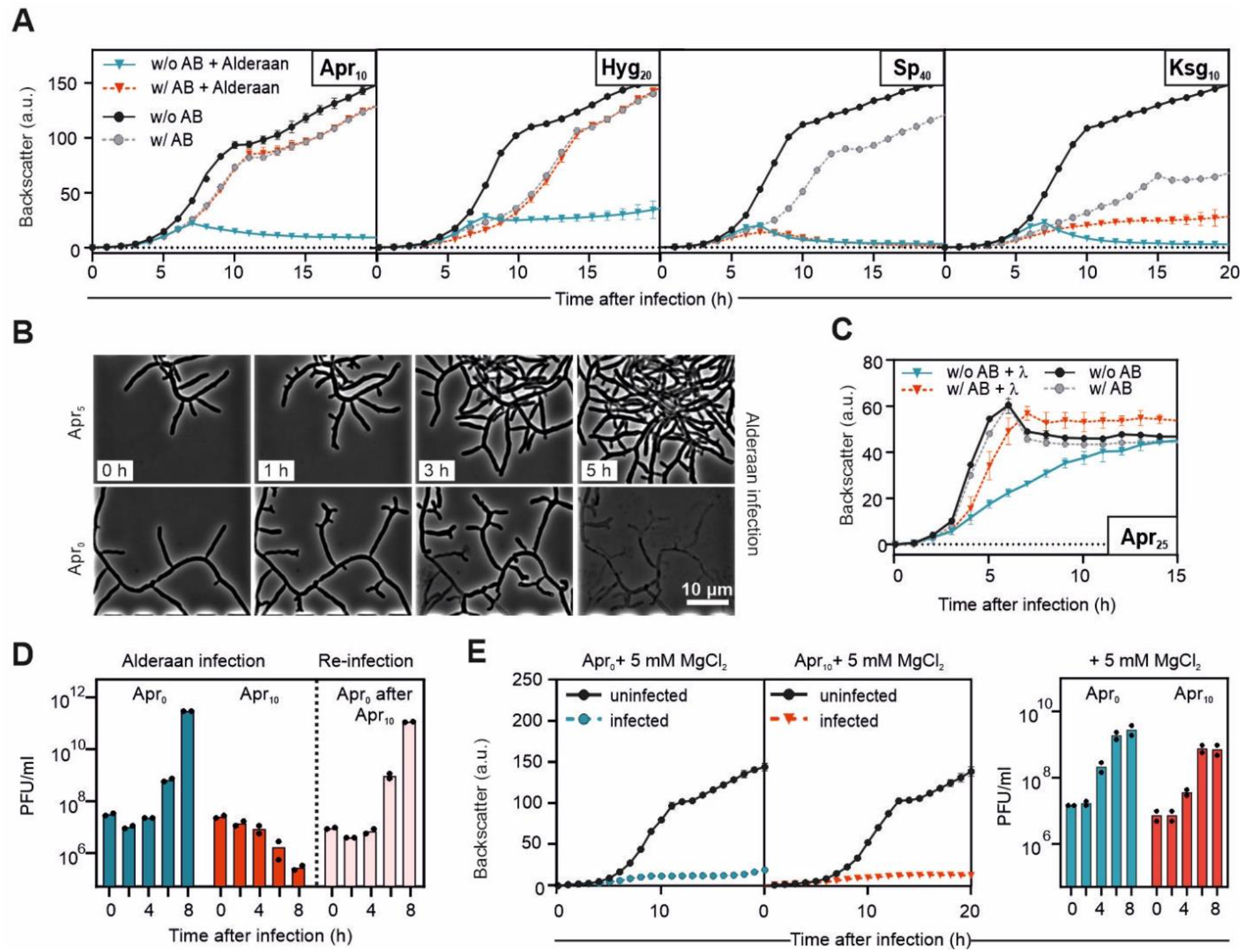
Aminoglycosides strongly inhibit phage amplification in liquid cultures. **A**, Infection curves for *Streptomyces venezuelae* infected by phage Alderaan in the presence of different aminoglycosides (concentrations, μg/ml indicated in subscript). **B**, Time-lapse micrographs of *S. venezuelae* cultivated in a microfluidics system and challenged with Alderaan (inserts: time after infection). **C**, Infection curves for *E. coli* infected by λ in presence of 25 μg/ml apramycin. **D,** Phage titers determined over two successive rounds of infection. A first infection round of *S. venezuelae* by Alderaan was performed in presence or absence of apramycin. At the end of the cultivation, surviving cells from the apramycin-treated cultures were collected and exposed to phage Alderaan again, this time in absence of apramycin. **E**, Effect of MgCl_2_ on infection of *S. venezuelae* by Alderaan, assessed by infection curves and determination of the corresponding phage titers over time. In **A**, **D** and **E**, Alderaan was added to an initial titer of 10^7^ PFU/ml; in **C**, λ was added to an initial titer of 10^8^ PFU/ml. For growth curves in **A**, **C** and **E**, data represent an average of three independent biological replicates (n=3); phage titers in **D** and **E** were quantified from two independent biological replicates (n=2).

Infection of the model phage λ was also strongly impaired in the presence of aminoglycosides. Here, concentrations as low as 25 μg/ml apramycin or kanamycin showed a protective effect in liquid cultures (Figure 2C and Supplementary Figure S2A) as well as an up to 1,000-fold reduction in plaque-forming units (Figure 1B and 1C). Furthermore, this effect was shown to be independent of the used host strain (Supplementary Figure S2B).

In the case of temperate phages such as λ, an increased entry in the lysogenic cycle could explain the absence of phage amplification in presence of aminoglycosides. To test this hypothesis, we conducted a re-infection experiment, in which cells surviving the first round of infection were washed and exposed to the same phage again. In the first infection round, cultures without apramycin showed a strongly increasing phage titer associated with extensive lysis of the culture. In contrast, infection in presence of apramycin was completely inhibited, showing no phage amplification during λ infection and even a ~100-fold decrease in phage titers over time for Alderaan (Figure 2D and Supplementary Figure S2C). Interestingly, removal of the antibiotic and re-infection of cells from apramycin-treated cultures resulted in similar amplification kinetics of Alderaan and λ as compared to an untreated control. Hence, these results do not support an increased formation of lysogens or resistance traits, but rather indicate a reversible antiphage effect of apramycin.

Since elevated Mg^2+^ levels were previously shown to interfere with aminoglycoside uptake^14^ and streptomycin-mediated inhibition of phage infection^12^, we examined whether the antiviral effect of apramycin is alleviated in the presence of MgCl_2_. As shown in Figure 2E, phage infection was completely restored by the addition of 5 mM MgCl_2_, as evidenced by the strong growth defect and the increasing phage titer during infection. Comparable results regarding the antagonistic effects of MgCl_2_ were also obtained for λ (Supplementary Figure S2D). Overall, these results suggest that the antiviral effect of aminoglycosides is based on an interference with phage infection at the intracellular level, probably during or shortly after phage DNA injection.

### Spent medium of natural aminoglycoside producers provides protection against phage predation

As *Streptomyces* are the natural producers of aminoglycosides, we examined whether infection of *S. venezuelae* in spent medium of the apramycin-producer *Streptomyces tenebrarius* provides protection against phage predation. Alderaan infection was not impaired by spent medium of *S. tenebrarius* harvested after one day of cultivation. In contrast, cultivation in spent medium taken after two days completely reproduced the antiviral effect observed during experiments with supplemented purified apramycin, showing equivalent growth of infected and uninfected cultures (Figure 3A). End-point quantification of extracellular phage titers confirmed this inhibition of infection, as no more extracellular phages were detectable in the supernatants of the infected cultures (Figure 3B). Importantly, this protective effect of *S. tenebrarius* spent medium coincided with the presence of apramycin in cultures, as determined by LC-MS (Figure 3C).

**Figure 3.**
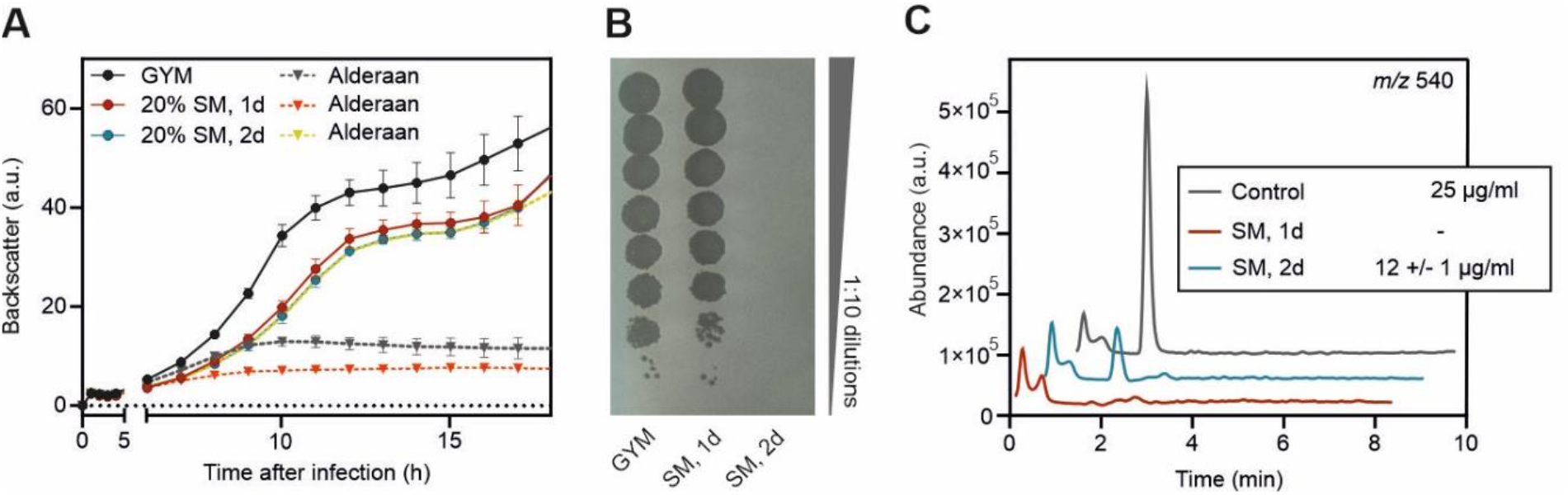
Secondary metabolites produced by *S. tenebrarius* inhibit phage infection. **A**, Influence of spent medium from *S. tenebrarius* on infection of *S. venezuelae* by Alderaan. Data represent an average of three independent biological replicates; error bars represent standard deviations. **B**, Determination of the final phage titers of infected cultures shown in **a**. Results are representative of two biological replicates. **C**, Extracted ion chromatogram of samples analysed by LC-MS assessing the presence of apramycin (molecular weight: 539.58 g/mol) in spent medium (SM) of *Streptomyces tenebrarius*. The indicated concentrations of apramycin are close to the detection limit under these measuring conditions. GYM: Glucose Yeast Malt extract medium.

Similar results could be obtained in experiments with spent medium of cultures of the natural kasugamycin-producer *Streptomyces kasugaensis.* However, this effect appears to be largely concentration-dependent. While supplementation with 4% of spent medium inhibited the phage-mediated culture collapse, showing the antiphage properties of *S. kasugaensis* supernatants (Figure S3), increased levels (20%) strongly delayed *S. venezuelae* growth, which is in line with the toxic effect of purified kasugamycin on growth of *S. venezuelae* (Figure 2A). Taken together, these data suggest that production of aminoglycoside antibiotics in natural environments might serve as a chemical defense providing protection against phage infection on a community level.

### Aminoglycosides block an early step of phage infection

To decipher the mechanism underlying the antiviral activity of aminoglycosides, we investigated the influence of apramycin on the different steps of the phage infection cycle (Figure 4A).

**Figure 4.**
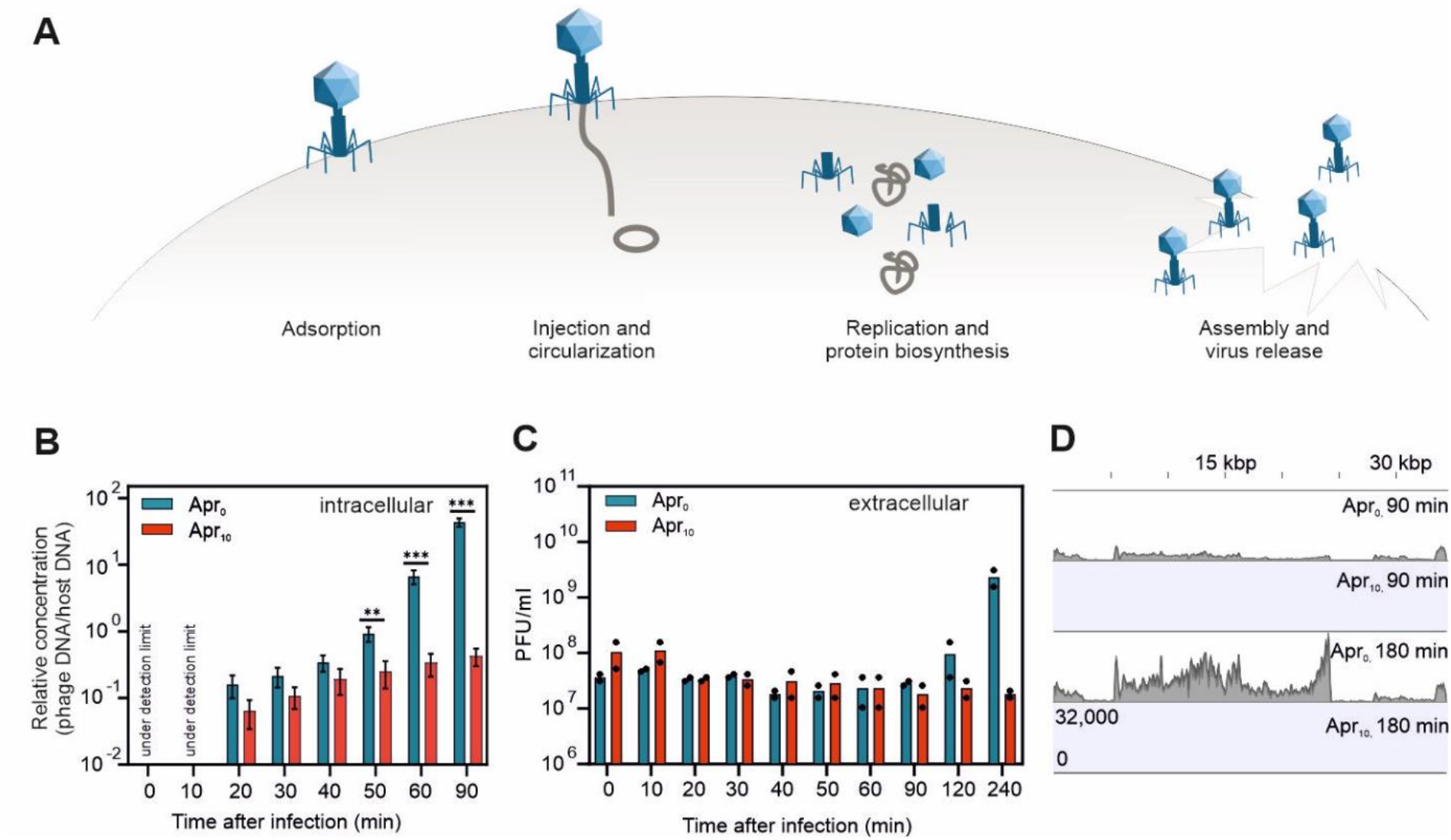
Apramycin blocks the phage life cycle at an early stage – before replication and transcription of phage DNA. **A**, Scheme of the phage lytic lifecycle, highlighting the different steps which could be inhibited by antiphage metabolites. **B**, Infection of *S. venezuelae* by Alderaan; time-resolved quantification of phage DNA by qPCR in the intracellular fraction. Data represent mean values of two independent biological replicates measured as technical duplicates. Significance was determined by using unpaired Student t-test (*** p<0.001; ** p<0.01). **C**, Time-resolved determination of Alderaan titers in the extracellular medium via double-agar overlays. n=2 independent replicates. **D**, RNA-seq coverage of the Alderaan genome (39 kbp) during infection in presence and absence of apramycin.

As a first approach, we assessed phage adsorption as well as DNA delivery and DNA amplification by determining the level of intra- and extracellular Alderaan DNA in the early steps of infection via quantitative real-time PCR. For both infection conditions, a comparable early infection phase within the first 40 min of infection was observed (Figure 4B, Figure S4A). Increasing intracellular phage DNA levels were in accordance with the simultaneously decreasing concentration of extracellular phage DNA shown in the adsorption assay (Supplementary Figure S4A). Interestingly, significant differences in intracellular phage DNA could be detected between 50-90 min after infection. In absence of apramycin, the exponential increase in intracellular phage DNA and the following increase in extracellular phage titers at 120 min after infection reflected the successful completion of the first infection cycles and the release of new virions. Conversely, only a slight, more linear increase in intracellular DNA and even a decrease in extracellular phage titers were obtained for infection under apramycin pressure. These results suggest an unaltered injection of phage DNA, but an inhibition of subsequent replication (Figure 4B and 4C). This inhibition of phage DNA replication was further evidenced by the long-term quantification of intracellular phage DNA during the entire infection. After eight hours, the level of intracellular phage DNA was about five orders of magnitude lower in the presence of apramycin in comparison to the untreated control (Supplementary Figure S4B).

Assuming that apramycin already blocks an early step of phage infection prior to genome replication, addition of the antibiotic after the replication phase would not interfere with the infection. This hypothesis was indeed confirmed by supplementation of the aminoglycoside at different time-points post-infection (Supplementary Figure S4C). Corresponding infection assays indicated that apramycin addition 30 min after infection was sufficient to prevent a reproductive Alderaan infection. The observed decrease in extracellular phage titers is probably the result of adsorption and following DNA injection of a fraction of phages without release of new infective viral particles.

In contrast, no decrease in extracellular phage titers was observed when adding apramycin one to two hours after infection, indicating that the first phages were able to complete their infection cycle before apramycin was added. Comparison of these results with the quantification of intracellular phage DNA (Figure 4B) further showed that this period corresponds to the replication phase, pointing out replication as an essential time point for the antiviral activity of aminoglycosides. In the case of the *E. coli* system, the measurement of potassium efflux is an established approach to probe the successful delivery of phage DNA into the bacterial cell^15^. Applying this method to infection of *E. coli* with phage λ confirmed that the injection process was not impaired by apramycin (Supplementary Figure S2E).

Next, we examined the influence of apramycin on phage DNA transcription. RNA-sequencing revealed an increasing transcription of Alderaan DNA during phage infection under normal infection conditions, whereas addition of apramycin prevented phage gene expression completely (Figure 4D and Figure S4D). In accordance with the previous results, these data suggest a blockage of phage infection prior to phage DNA replication and transcription, which is congruent with a recent report of inhibition of two mycobacteriophages by streptomycin, kanamycin and hygromycin^16^.

To visualize intracellular phage infections in the presence and absence of apramycin, we performed fluorescence *in situ* hybridization of phage DNA (phage targeting direct-geneFISH) using Alexa647-labelled probes specific for the particular phage genome. For λ, similar amounts of injected phage DNA were detected for both infection conditions after 30 min. This result is in line with the potassium efflux assay described above which showed similar injection kinetics in the presence of apramycin (Figure S2E). As the infection progressed, only samples without apramycin exhibited an increase in fluorescence intensity 90 min post-infection, further hinting at an inhibited replication in the presence of apramycin (Figure 5A and 5B). In this assay, the formation of bright and distinct fluorescent foci is indicative for advanced viral infections^17^. For Alderaan, an increase in red fluorescence and thus intracellular phage DNA could be observed 4 hours after infection, reflecting DNA replication. In contrast, apramycin-treated samples showed only a weak and more diffuse red fluorescent signal in the 8 h samples (Figure 5C). Quantifying the relative background-corrected total intensities at 8 h after infection indicated a markedly increased intensity for samples without apramycin in comparison to apramycin-treated samples (Figure 5D), which was in line with the results obtained for λ infection. Signals arising from single phage genomes (right after injection) were not detected at earlier time points. However, this signal might be obscured by the strong autofluorescence of *Streptomyces*.

**Figure 5.**
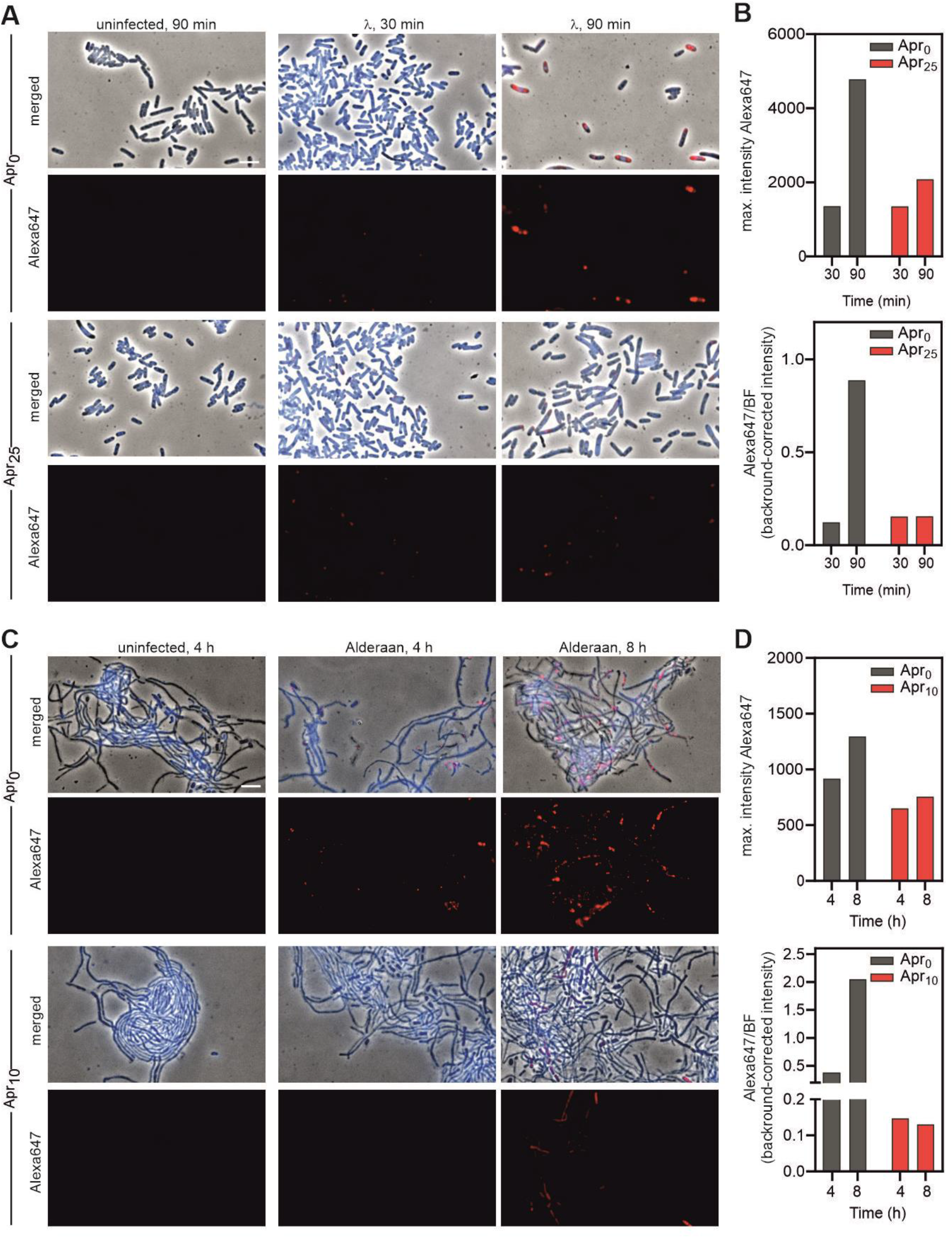
Visualization of intracellular phage DNA by phage targeting direct-geneFISH. **A** and **C,** Phage targeting direct-geneFISH micrographs of (**A**) *E. coli* infected with λ and (**C**) *S. venezuelae* infected with Alderaan in presence and absence of 25 μg/ml and 10 μg/ml apramycin, respectively. First and third row: phase-contrast pictures merged with fluorescence signal from bacterial DNA (DAPI, blue) and phage DNA (Alexa647, red). Second and fourth row: Fluorescence signal from phage DNA only (Alexa647, red). Scale bar: 5 μm. **B** and **D**, Quantification of Alexa647 fluorescence in (**B**) *E. coli* cells infected with λ and (**D**) *S. venezuelae* cells infected with Alderaan, shown as the maximal and background-corrected intensity of fluorescent signals.

### Acetylation of apramycin abolishes its antibacterial, but not antiphage properties

Enzymatic modification of aminoglycosides is a major mechanism of bacterial resistance to these antibiotics. Aminoglycoside-modifying enzymes are categorized in three major classes: aminoglycoside *N*-acetyltransferases (AACs), aminoglycoside *O*-nucleotidyltransferases (ANTs) and aminoglycoside *O*-phosphotransferases (APHs)^18^. Addition of an acetyl, adenyl or phosphoryl group at various positions of the aminoglycoside core scaffold decreases the binding affinity of the drug for its primary ribosomal target, leading to the loss of the antibacterial potency - the modified aminoglycosides being described as “inactivated”. However, the impact of these modifications on the antiphage activity of aminoglycosides is unknown. We set out to answer this question using apramycin and the acetyltransferase AAC(3)IV. In presence of apramycin, AAC(3)IV catalyzes the acetylation of the 3-amino group of the deoxystreptamine ring, using acetyl-CoA as a co-substrate (Figure 6A).

Using purified AAC(3)IV, we performed an *in vitro* acetylation reaction of apramycin. LC-MS analysis of the reaction mixtures revealed a complete acetylation of apramycin, as the peak of apramycin (*m/z* 540) disappeared in favor of the one corresponding to acetylated apramycin (*m/z* 582) (Figure 6B).

**Figure 6.**
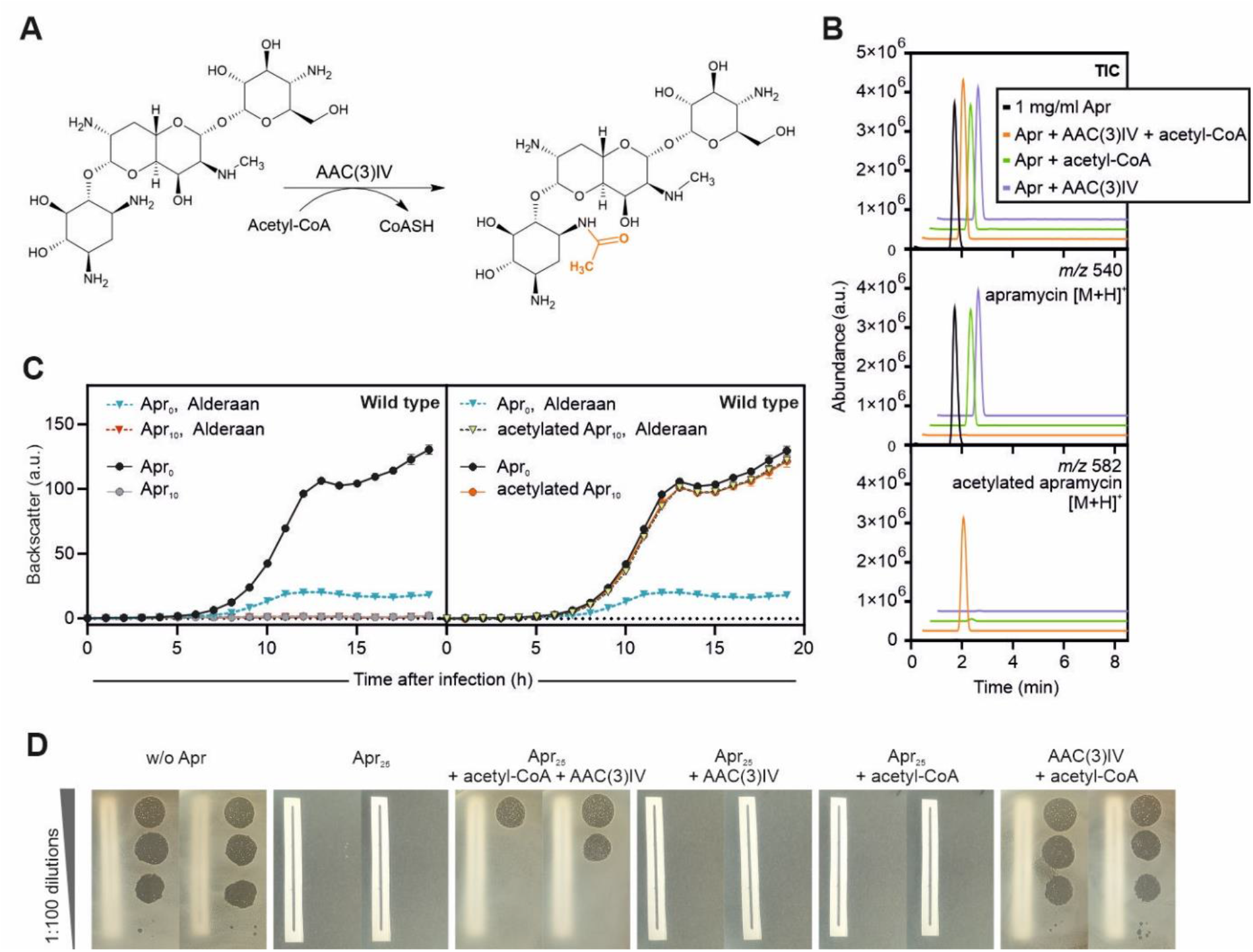
Acetylated apramycin strongly inhibits phage infection, despite the loss of its antibacterial properties. **A,** Acetylation reaction of apramycin catalyzed by the AAC(3)IV acetyltransferase**. B,** Total ion chromatogram (TIC) and extracted ion chromatograms of samples analysed by LC-MS assessing the presence of apramycin (molecular weight: 539.58 g/mol, *m/z* 540) and acetylated apramycin (molecular weight: 581.62 g/mol, *m/z* 582) after *in vitro* acetylation of apramycin. **C**, and **D**, Effect of acetylated apramycin on infection of wild type *S. venezuelae* with Alderaan, performed in liquid (**C**) and solid medium (**D**). In **D**, a piece of paper was placed below plates to facilitate assessment of bacterial growth.

The efficiency of the acetylation reaction being confirmed, we tested the effect of acetylated apramycin on phage infection in liquid medium, using wild-type *S. venezuelae* (not encoding a plasmid-borne acetyltransferase) and its phage Alderaan. As expected, apramycin fully prevented growth of *S. venezuelae*, while acetylated apramycin did show any toxicity effect. Strikingly, phage infection was completely inhibited in presence of acetylated apramycin, suggesting that acetylation of apramycin does not interfere with its antiphage properties (Figure 6C). Plate assays showed a comparable pattern: acetylation of apramycin suppressed its antibacterial effect, but did not disrupt its ability to inhibit phage infection (Figure 6D). Altogether, these results suggest a decoupling of the antibacterial and antiphage properties of apramycin and further highlight the distinct molecular target accounting for apramycin’s antiphage properties.

## Discussion

We have shown that aminoglycosides inhibit phage infection in a diverse set of bacterial hosts by blocking an early step of the phage life cycle between DNA injection and replication. These findings highlight the multifunctionality of this class of antibiotics, as they possess both antibacterial and antiviral properties.

The versatility of aminoglycosides can be attributed to their ability to bind a wide variety of nucleic acids - DNA or RNA, biologically or non-biologically derived. The most prominent target of aminoglycosides is the 16S rRNA, accounting for the disruption of protein translation and hence their bactericidal properties^18^. Aminoglycosides have also been shown to bind to seemingly unrelated families of RNA molecules such as group I introns^19^, a hammerhead ribozyme^20^, the *trans*-activating response element (TAR)^21^ and the Rev response element (RRE) of the human immunodeficiency virus HIV^22–24^. Furthermore, *in vitro* studies showed condensation of purified phage λ DNA by aminoglycosides. It was proposed that the clamp formed by aminoglycosides around the DNA double helix causes a bend responsible for the formation of toroids and other structural deformations^25,26^.

Injected phage DNA is linear, in a relaxed state and not protected by DNA-binding proteins, and therefore probably highly sensitive to DNA-binding molecules. Interestingly, anthracyclines – another class of secondary metabolites produced by *Streptomyces* with antiphage properties – inhibit phage infection at a similar stage^5^. While the exact mechanism of action underlying phage inhibition by anthracyclines and aminoglycosides remains elusive, these recent results suggest that already injected but not yet replicating phage DNA is preferentially targeted by antiviral molecules.

Therapeutical use of phages - known as phage therapy - is often combined with an antibiotic treatment due to the potentially synergistic effect between these two antimicrobial agents. In contrast, we describe here an antagonistic impact of a common antibiotic class on phages, which has important implications for phage-aminoglycoside combination treatment. We propose that sensitivity of the phage to aminoglycosides should be assessed *in vitro* before administration of such combination therapy.

From a more fundamental perspective, these findings also shed new light on the role of aminoglycosides in natural bacterial communities. While their use as antibiotics for medical applications has been extensively documented, until now relatively little was known about their function in the natural setting. We posit that aminoglycosides are not only used by their producers as a powerful weapon against bacterial competitors, but also protect them against phage predation at the community level. Considering the colossal number of molecules produced by environmental bacteria whose physiological role is still unclear, we postulate that additional prokaryotic antiphage metabolites are to be discovered in the future, further underlining the extraordinary diversity of strategies employed by bacteria against their viral predators.

## Material and methods

### Bacterial strains and growth conditions

All bacterial strains, phages and plasmids used in this study are listed in Supplementary Tables S2, S3 and S4, respectively. For growth studies and double-agar overlay assays, *Streptomyces spp*. cultures were inoculated from spore stocks and cultivated at 30 °C and 120 rpm using Glucose Yeast Malt extract (GYM) medium for *S. venezuelae, S. tenebrarius* and *S. kasugaensis;* and Yeast Extract Malt Extract (YEME) medium for *S. coelicolor*^27^. Cultivation of *E. coli* was performed in lysogeny broth (LB) medium at 37 °C and 170 rpm, while *C. glutamicum* was grown in brain heart infusion (BHI) medium at 30°C and 120 rpm.

For double agar overlays, BHI agar for *C. glutamicum*, LB agar for *E. coli* and GYM agar (pH 7.3) for all *Streptomyces* species was used, with 0.4% and 1.5% agar for the top and bottom layers, respectively. For quantification of extracellular phages, 2 μl of the culture supernatants were spotted on a bacterial lawn propagated on a double-agar overlay inoculated at an initial OD_450_=0.4 for *Streptomyces* spp., OD_600_=0.1 for *E. coli* and OD_600_= 0.7 for *C. glutamicum*. Both agar layers were supplemented with antibiotics at the indicated concentrations.

For standard cloning applications, *E. coli* DH5α was cultivated in LB medium containing the respective antibiotic at 37°C and 120 rpm. For conjugation between *Streptomyces spp.* and *E. coli*, the conjugative *E. coli* ET12567/pUZ8002 strain was used^28^.

### Recombinant DNA work and cloning

All plasmids and oligonucleotides used in this study are listed in Supplementary Tables S4 and S5, respectively. Standard cloning techniques such as PCR and restriction digestion were performed according to standard protocols^29^. In all cases, Gibson assembly was used for plasmid construction^30^. DNA regions of interest were amplified via PCR using the indicated plasmid DNA as template. The plasmid backbone was cut using the listed restriction enzymes.

DNA sequencing and synthesis of oligonucleotides was performed by Eurofins Genomics (Ebersberg, Germany).

### Phage infection curves

For phage infection curves, the BioLector microcultivation system of m2p-labs (Aachen, Germany) was used^31^. Cultivations were performed as biological triplicates in FlowerPlates (m2plabs, Germany) at 30 °C and a shaking frequency of 1200 rpm. During cultivation, biomass was measured as a function of backscattered light intensity with an excitation wavelength of 620 nm (filter module: λ_Ex_/ λ_Em_: 620 nm/ 620 nm, gain: 25) every 15 minutes. Main cultures of *Streptomyces spp*. in 1 ml GYM medium containing the indicated supplements were inoculated with overnight cultures in the same medium to an initial OD_450_= 0.15. Infection was performed by adding phages to an initial phage titer of 10^7^ PFU/ml. Supernatants were collected in 2 h intervals to determine the time course of phage titer via double agar overlays. Phage infection curves in *E. coli* were done in the same way at 37°C and 1200 rpm using an initial OD_600_=0.1 in 1 ml LB medium and an initial phage titer of 10^8^ PFU/ml, resulting in a multiplicity of infection (MOI) of 1.

Phage infection curves in shaking flasks were performed analogously to the cultivation in microbioreactors using a shaking frequency of 120 rpm.

### Cultivation and perfusion in microfluidic device

Single-cell analysis of *S. venezuelae* cells infected with phage Alderaan in presence and absence of apramycin was performed using an in-house developed microfluidic platform ^32–34^. Cultivation and time-lapse imaging were performed in three steps. First, cultivation chambers in the microfluidic chip were inoculated with GYM medium containing an initial spore titer of 10^8^ PFU/ml. During the following pre-cultivation phase, cells in all chambers were cultivated under continuous GYM medium supply supplemented with 2.5 μg/ml apramycin (flow rate: 300 nl/min) to allow comparable growth conditions. After 6 h of pre-cultivation, cells were cultivated for 3 h in GYM medium containing one of the final apramycin concentrations (0, 5 or 10 μg/ml). Subsequently, infection was initiated by continuous supply of GYM medium containing the final apramycin concentrations and Alderaan phages with a titer of 10^8^ PFU/ml (flow rate 200 nl/min). By using disposable syringes (Omnifix-F Tuberculin, 1 ml; B. Braun Melsungen AG, Melsungen, Germany) and high-precision syringe pump system (neMESYS, Cetoni GmbH, Korbussen, Germany) continuous medium supply and waste removal was achieved. Phase contrast images were obtained in 5 min intervals (exposure time 100 ms) by a fully motorized inverted Nikon Eclipse Ti microscope (Nikon GmbH, Düsseldorf, Germany). During the complete cultivation, the temperature was set to 30°C using an incubator system (PeCon GmbH, Erbach, Germany).

### Cultivation in spent medium

For preparation of spent medium, cultures of the natural apramycin producer *Streptomyces tenebrarius* were prepared by inoculating 50 ml of GYM medium to an initial OD_450_ of 0.1 and were cultivated for 4 days. Spent medium of the culture was collected every day by centrifugation and subsequent filtration of the supernatant. After adjusting the pH to 7.3, GYM medium and spent medium were mixed in a ratio of 4:1—so that spent medium accounts for 20% of the total volume. Ten-times concentrated GYM was added to keep the concentration of C-sources equal to the one of fresh GYM medium. Cultivation and infection of the apramycin-resistant *S. venezuelae* – pIJLK04 strain in 20% spent medium was conducted in microbioreactors as described previously by using an initial OD_450_ = 0.5 and an initial phage titer of 10^6^ PFU/ml.

Preparation of spent medium of the natural kasugamycin producer *S. kasugensis* and subsequent cultivation and infection of the kasugamycin-resistant *S. venezuelae* – pIJLK03 in 4% and 20% spent medium was performed analogously.

### LC-MS measurements of apramycin

Aminoglycosides were analysed using an Agilent ultra-high-performance LC (uHPLC) 1290 Infinity System coupled to a 6130 Quadrupole LC-MS System (Agilent Technologies, Waldbronn, Germany). LC separation was carried out using an InfinityLab Poroshell 120 2.7 μm EC-C_18_ column (3.0 × 150 mm; Agilent Technologies, Waldbronn, Germany) at 40 °C. For elution, 0.1 % acetic acid (solvent A) and acetonitrile supplemented with 0.1 % acetic acid (solvent B) were applied as the mobile phases at a flow rate of 0.3 ml/min. A gradient elution was used, where the amount of solvent B was increased stepwise: minute 0 to 6: 10 % to 25 %, minute 6 to 7: 25 % to 50 %, minute 7 to 8: 50 % to 100 % and minute 8 to 8.5: 100 % to 10 %. The mass spectrometer was operated in the positive electrospray ionization (ESI) mode, and data were acquired using the selected-ion-monitoring (SIM) mode. An authentic apramycin standard was obtained from Sigma-Aldrich (Munich, Germany). Area values for [M+H]^+^ mass signals were linear for metabolite concentrations from 10 to 50 μg/ml.

### Potassium efflux assays

Cultures of *E. coli* LE392 - pEKEx2.d were grown in LB medium supplemented with 50 μg/ml apramycin at 37 °C and 170 rpm overnight. Fresh LB medium (50 μg/ml apramycin if needed) was inoculated 1:100 from the overnight cultures and incubated at 37 °C and 120 rpm for 1.5 h. The cultures were centrifuged at 5000 g for 20 min and the pellets were resuspended in SM buffer (0.1 M NaCl, 8 mM MgSO_4_, 50 mM Tris-HCl, pH 7.5), OD_600_ was measured and adjusted to OD_600_ = 2. The cultures were stored at 4 °C and incubated at 37 °C for 5 min directly before use. The measurements were performed using an Orion potassium ion selective electrode (Thermo Scientific). 5 ml of the prepared cultures were mixed 1:50 with ionic strength adjuster (ISA, Thermo Scientific) and measurements were started immediately monitoring the electric potential in mV every 5 s for a total of 60 min at room temperature under constant stirring. If apramycin was needed, it was added in the beginning to a concentration of 100 μg/ml. After 5.5 min, 100 μl of a PEG-precipitated λ phage lysate in SM buffer (10^11^ PFU/ml) was added to the cultures.

### Quantitative real-time PCR

Quantification of cell-associated Alderaan phages was performed via quantitative real-time PCR. For this, infection was performed as described in ‘Phage infection curves‘. At the indicated time points, 2 OD units of cells were harvested via centrifugation at 5,000 *g* and 4 °C for 15 min and washed two times with PBS. Cell disruption was subsequently performed by resuspending the cell pellet in 150 μl PBS and incubation at 95 °C and 1000 rpm for 15 min. After centrifugation at 16,000 g and 4°C for 10 min, 5 μl of the supernatant as template DNA was mixed with 10 μl 2x Universal qPCR Master Mix (New England BioLabs) and 1 μl of each oligonucleotide (Table S5, final oligonucleotide concentration: 0.5 μM) and adjusted to a final volume of 20 μl with dd. H_2_O. Measurements were performed in 96-well plates in the qTOWER 2.2 (Analytik Jena). For the determination of the relative concentration of cell-associated phages, the relative expression ratio of the phage target phage gene (coding for the minor tail protein of Alderaan) to the housekeeping gene *atpD* (coding for ATP synthase beta subunit) was calculated via ‘Relative quantification method’ of the qPCRsoft 3.1 software (Analytik Jena).

### Transcriptomics via RNA-sequencing

To compare transcription of phage and host DNA in presence and absence of apramycin, infection of the apramycin-resistant strain *S. venezuelae* ATCC 10712 - pIJLK04 with Alderaan was conducted as described in ‘Phage infection curves‘. Cells were harvested 90 min and 180 min after infection. RNA purification was done using the Monarch Total RNA Miniprep Kit (New England Biolabs) according to the manufacturer’s manual. Depletion of rRNA, library preparation and sequencing were conducted by GENEWIZ (Leipzig, Germany).

After sequencing, all subsequent steps were conducted using CLC genomic workbench V.20.0.4 software (QIAGEN, Hilden, Germany). The initial quality check to analyze read quality and sequencing performances was followed by a trimming step. This step was used to remove read-through adapter sequences, left-over adapter sequences, low quality reads (limit = 0.05) and ambiguous nucleotides. Subsequently, the trimmed reads were mapped against the genomes of *S. venezuelae* (accession no.: NC_018750.1) and the phage Alderaan (accession no.: MT711975.1). Coverage plots were generated to show the distribution of mapped reads on both genomes. Subsequently, transcripts per million (TPM) values were calculated using the RNA-seq analysis tool of CLC genomics workbench (read alignment parameters: Mismatch cost: 2, Insertion cost: 3, Deletion cost: 3, Length fraction: 0.8, Similarity fraction: 0.8, Strand specific: both, Maximum number of hits for a read: 10). A table containing these values as well as an overview matrix containing all values were exported for each sample. Raw data as well as processed tables were deposited in the GEO database under the accession number GSE171784.

### Phage targeting direct-geneFISH

Visualization and quantification of intracellular phage DNA during the time course of infection was conducted via fluorescence *in situ* hybridization (FISH), following the direct-geneFISH protocol^35^, with modifications as described below.

Design of phage gene probes was done using the gene-PROBER^36^. Sequences of the 200 bp polynucleotides for Alderaan and 300 bp polynucleotides for λ are provided in Supplementary Table S6. Phage infection was performed as described in the ‘Phage infection curves’ section. For infection of *E. coli*, the chemical labelling of polynucleotides with Alexa Fluor 647 dye (Thermo Fisher Scientific) as well as the “core” direct-geneFISH protocol for microscopic slides was conducted as described previously^35^. Imaging of cells was performed with an inverted time-lapse live cell microscope (Nikon TI-Eclipse, Nikon Instruments, Germany) using a 1003 oil immersion objective (CFI Plan Apo Lambda DM 100X, NA 1.45, Nikon Instruments, Germany)^32^. Fluorescence was recorded using the optical filters DAPI-U HQ and CY5 HQ (DAPI: excitation 395/25, dichroic: 425, emission: 460/50, exposure time: 500 ms; Cy5: excitation 620/60, dichroic: 660, emission: 700/75, exposure time: 500 ms (Nikon TI-Eclipse, Nikon Instruments, Germany)). Phase contrast was imaged with an exposure time of 200 ms.

For *S. venezuelae* infection the protocol was adjusted as follows: Fixation of cells and phages was performed in 50 % ethanol overnight. After washing and immobilization, permeabilization was performed in two steps using 1.25 mg/ml lysozyme for 60 min at 37 °C and subsequently 60 U/ml Achromopeptidase for 15 min at 37°C. Due to the high GC-content of phage Alderaan, formamide concentration in the hybridization buffer and in the humidity chamber was adjusted to 60% and NaCl concentration in the washing buffer was reduced to 14 mM NaCl. After counterstaining with DAPI, imaging of cells was performed as described for *E. coli* using the optical filters DAPI-U HQ and CY5 HQ with the indicated exposure times (DAPI: excitation 395/25, dichroic: 425, emission: 460/50, exposure time: 600 ms; Cy5: excitation 620/60, dichroic: 660, emission: 700/75, exposure time: 800 ms (Nikon TI-Eclipse, Nikon Instruments, Germany)). Phase contrast was imaged with an exposure time of 200 ms. The images for phage signal quantification were taken at the same exposure times to enable comparison; exposure times were adjusted so as to avoid overexposure of the signals.

For quantification of the microscopic analyses, the relative background-corrected total intensities (rel. Intbg) of each sample were calculated. To this end, the total intensities of the Cy5 image as well as of the inverted bright field (BF) images were measured. Subsequently, three representative areas of the background were chosen for each image to be used as a correction factor of the total intensities (bgint). These background correction factors were multiplied with the whole image area (area) and subtracted from the total intensities (Int) of the corresponding channel. In a final step, the background-corrected intensity of Cy5 was normalized with regard to the background-corrected intensity of BF. The resulting rel. Intbg represents the total phage signal per cell intensity.

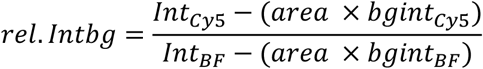

### Purification of the AAC(3)IV apramycin acetyltransferase

For heterologous protein overproduction, *E. coli* BL21 (DE3) cells containing the pAN6_aac(3)IV_CStrep plasmid were cultivated as described in bacterial strains and growth conditions. Pre-cultivation was performed in LB medium supplemented with 50 μg/ml kanamycin (LB Kan_50_), which was incubated overnight at 37 °C and 120 rpm. The main culture in LB Kan_50_ medium was inoculated to an OD_600_ of 0.1 using the pre-culture. At an OD_600_ of 0.6 gene expression was induced using 100 μM IPTG. Cells were harvested after additional 24 h incubation at 20 °C.

Cell harvesting and disruption were performed as described earlier^37^ using buffer A (100 mM Tris-HCl, pH 8.0) for cell disruption and buffer B (100 mM Tris-HCl, 500 mM NaCl, pH 8.0) for purification. Purification of the Strep-tagged AAC(3)IV apramycin acetyltransferase was conducted by applying the supernatant to an equilibrated 2 ml Strep-Tactin-Sepharose column (IBA, Göttingen, Germany). After washing with 20 ml buffer B, the protein was eluted with 5 ml buffer B containing 15 mM D-desthiobiotin (Sigma Aldrich).

After purification, the purity of the elution fractions was checked by SDS-PAGE^38^ using a 4– 20% Mini-PROTEAN gradient gel (Bio Rad, Munich, Germany). Protein concentration of the elution fraction was determined with the Pierce BCA Protein Assay Kit (ThermoFisher Scientific, Waltham, MA, USA) and the elution fraction with the highest protein concentration was chosen for further use.

### In vitro acetylation reaction of apramycin

Protein purification of the AAC(3)IV apramycin acetyltransferase was conducted as described above. Acetylation of apramycin was performed using a modified version of the protocol from Magalhaes and Blanchard^39^. Assay mixtures were composed of 100 μl 100 mM Tris-HCl, 500 mM NaCl, pH 8.0 containing the AAC(3)IV at a concentration of 10 μg/ml, as well as 10 mM apramycin (approximately 5 mg/ml) and 10 mM acetyl-CoA sodium salt (Sigma Aldrich). The assay mixtures were incubated at 37°C for 20 minutes.

## Acknowledgements

We thank the European Research Council (ERC Starting Grant, grant number 757563) and the Helmholtz Association (grant number W2/W3-096) for financial support.

We thank Paul Ramp and Natalia Tschowri for providing strains and plasmids, and our bachelor student Lisa Helm for her contribution to this project. We furthermore thank Mark Buttner (JIC, Norwich) for introducing us into *Streptomyces* biology and for many fruitful discussions.

## Author contributions

Conceptualization: A.H., L.K., J.F. | Data curation: M.H. and L.K. | Formal analysis: all | Funding acquisition: J.F. | Investigation: L.K., A.H., T.L., C.G., M.H., L.M., J.Wie. | Methodology: L.K., A.H., T.L., M.H., C.M., J.Wie., L.M., J.F. | Project administration: J.F. | Resources: J.Wit. | Supervision: J.F. | Validation: A.H., L.K., T.L., M.H., L.M., J.M., J.F. | Visualization: L.K., A.H., J.F. | Writing – original draft: A.H., L.K., J.F. | Writing – review and editing: all

## Supplementary information

### Tables

Table S1 | Aminoglycoside-modifying enzymes used in this study

Table S2 | Bacterial strains used in this study.

Table S3 | Phages used in this study

Table S4 | Plasmids used in this study

Table S5 | Oligonucleotides used in this study

Table S6 | Polynucleotides used for phage targeting direct-geneFISH

### Figures

Figure S1 | Dose-dependent effect of apramycin on *Streptomyces* phage Alderaan

Figure S2 | Effect of aminoglycosides on *E. coli* phage λ.

Figure S3 | Secondary metabolites produced by *Streptomyces kasugaensis* inhibit phage infection.

Figure S4 | Investigations of the mechanism of action of apramycin.

### Videos

Video S1: Apramycin prevents cell lysis during infection of *S. venezuelae* with phage Alderaan.

