## Supplemental Information for "Aminoglycoside antibiotics inhibit phage infection by blocking an early step of the phage infection cycle"

Running title: Aminoglycosides inhibit phage infection


Larissa Kever[1#], Aël Hardy[1#], Tom Luthe[1], Max Hünnefeld[1], Cornelia Gätgens[1], Lars Milke[1], Johanna Wiechert[1], Johannes Wittmann[2], Cristina Moraru[3], Jan Marienhagen[1,4] and Julia Frunzke[1*]

[1]Institute of Bio- und Geosciences, IBG-1: Biotechnology, Forschungszentrum Jülich, 52425 Jülich, Germany

[2]Leibniz Institute DSMZ—German Collection of Microorganisms and Cell Cultures, Inhoffenstraße 7B, 38124 Braunschweig, Germany

[3]Institute for Chemistry and Biology of the Marine Environment (ICBM), Carl-von-Ossietzky-University Oldenburg, Oldenburg, Germany

[4]Institute of Biotechnology, RWTH Aachen University, Aachen, Germany

[#] Authors contributed equally to this work.

*Corresponding author:

**Tables**

Table S1 | Aminoglycoside-modifying enzymes used in this study

Table S2 | Bacterial strains used in this study.

Table S3 | Phages used in this study

Table S4 | Plasmids used in this study

Table S5 | Oligonucleotides used in this study

Table S6 | Polynucleotides used for phage targeting direct-geneFISH

**Figures**

Figure S1 | Dose-dependent effect of apramycin on the *Streptomyces* phage Alderaan.

Figure S2 | Effect of aminoglycosides on *E. coli* phage λ.

Figure S3 | Secondary metabolites produced by *Streptomyces kasugaensis* inhibit phage infection.

Figure S4 | Investigations of the mechanism of action of apramycin.

**Videos**

Video S1: Apramycin prevents cell lysis during infection of *S. venezuelae* with phage Alderaan.

### Tables

**Supplementary Table S1: Aminoglycoside-modifying enzymes used in this study**

| Antibiotic | Gene | Annotation | Modification |
|---|---|---|---|
| **Apramycin** | *aac(3)IV* | Aminoglycoside *N*(3)-acetyltransferase | Acetylation of 3-amino group of the deoxystreptamine ring |
| **Hygromycin** | *aph(7'')-Ia* | Aminoglycoside *O*-phosphotransferase APH(7'')-Ia, | Phosphorylation of hydroxyl group at position 7'' |
| **Kanamycin** | *aphA1* | Aminoglycoside 3'-phosphotransferase | Phosphorylation of hydroxyl group at position 3‘ |
| **Kasugamycin** | *kac338* | Kasugamycin acetyltransferase | Acetylation of the 2'-$NH_2$ |
| **Spectinomycin/ Streptomycin** | *aadA* | Aminoglycoside (3'') (9) adenylyltransferase | *O*-adenylation at positions 3″ and 9 |

**Supplementary Table S2: Bacterial strains used in this study**

| Strains | Genotype | Reference |
|---|---|---|
| *C. glutamicum* MB001 | ATCC 13032 strain with deletion of prophages ΔCGP1 (cg1507-cg1524), ΔCGP2 (cg1746-cg1752) und ΔCGP3 (cg1890-cg2071) | [1] |
| *C. glutamicum* MB001 (DE3) | MB001 derivative with chromosomally encoded T7 gene *1* (cg1122-$P_{\mathit{lacI}}$-*lacI*-$P_{\mathit{lac}\mathrm{UV5}}$ –*lacZ*α-T7 gene *1*-cg1121) | [2] |
| *C. glutamicum* MB001 – pEKEx2a | MB001 carrying the plasmid pEKEx2a, $Kan^R$ | This study |
| *C. glutamicum* MB001 – pEKEx2b | MB001 carrying the plasmid pEKEx2b, $Hyg^R$ | This study |
| *C. glutamicum* MB001 (DE3) – pEKEx2c | MB001 (DE3) carrying the plasmid pMKEx-Kac, $Kan^R$ | This study |
| *C. glutamicum* MB001 – pEKEx2d | MB001 carrying the plasmid pEKEx2d, $Apr^R$ | This study |
| *C. glutamicum* MB001 – pEKEx2e | MB001 carrying the plasmid pEKEx2e, $Sp^R$/$Sm^R$ | This study |
| *Escherichia coli* DH5α | *supE44 ΔlacU169 (f80lacZ*DM15*) hsdR17 recA1 endA1 gyrA96 thi-1 relA1* | Invitrogen |
| *Escherichia coli* ET12567/pUZ8002 | *dam, dcm, hsdS, cat, tet*; carrying plasmid pUZ8002 | [3] |
| *Escherichia coli* DSM 613 | Wild-type strain | [4] |
| *E. coli* DSM 613 – pEKEx2a | *E. coli* DSM 613 carrying the plasmid pEKEx2a, $Kan^R$ | This study |
| *E. coli* DSM 613 – pEKEx2b | *E. coli* DSM 613 carrying the plasmid pEKEx2b, $Hyg^R$ | This study |
| *E. coli* DSM 613 – pEKEx2c | *E. coli* DSM 613 carrying the plasmid pEKEx2c, $Ksg^R$ | This study |
| *E. coli* DSM 613 – pEKEx2d | *E. coli* DSM 613 carrying the plasmid pEKEx2d, $Apr^R$ | This study |
| *E. coli* DSM 613 – pEKEx2e | *E. coli* DSM 613 carrying the plasmid pEKEx2e, $Sp^R$/$Sm^R$ | This study |
| *Escherichia coli* DSM 5695 | *$F^+$ met str $T1^s$ $T6^s$ $lambda^-$* | [5] |
| *E. coli* DSM 5695 – pEKEx2a | *E. coli* DSM 5695 carrying the plasmid pEKEx2a, $Kan^R$ | This study |
| *E. coli* DSM 5695 – pEKEx2b | *E. coli* DSM 5695 carrying the plasmid pEKEx2b, $Hyg^R$ | This study |
| *E. coli* DSM 5695 – pEKEx2c | *E. coli* DSM 5695 carrying the plasmid pEKEx2c, $Ksg^R$ | This study |
| *E. coli* DSM 5695 – pEKEx2d | *E. coli* DSM 5695 carrying the plasmid pEKEx2d, $Apr^R$ | This study |
| *E. coli* DSM 5695 – pEKEx2e | *E. coli* DSM 5695 carrying the plasmid pEKEx2e, $Sp^R$/$Sm^R$ | This study |
| *Escherichia coli* DSM 4230 | *$F^-$ hsdR514 ($rk^-$ $mk^-$) supE44 supF58 Δ(lacIZY)6 galK2 galT22 metB1 trpR55 $lambda^-$* | [6] |

| | | |
|---|---|---|
| *E. coli* DSM 4230 – pEKEx2a | *E. coli* DSM 4230 carrying the plasmid pEKEx2a, $Kan^R$ | This study |
| *E. coli* DSM 4230 – pEKEx2b | *E. coli* DSM 4230 carrying the plasmid pEKEx2b, $Hyg^R$ | This study |
| *E. coli* DSM 4230 – pEKEx2c | *E. coli* DSM 4230 carrying the plasmid pEKEx2c, $Ksg^R$ | This study |
| *E. coli* DSM 4230 – pEKEx2d | *E. coli* DSM 4230 carrying the plasmid pEKEx2d, $Apr^R$ | This study |
| *E. coli* DSM 4230 – pEKEx2e | *E. coli* DSM 4230 carrying the plasmid pEKEx2e, $Sp^R$/$Sm^R$ | This study |
| *Escherichia coli* JW3996 | *E. coli* BW25113 Δ*lamB* | [7] |
| *Streptomyces venezuelae* ATCC 10712 | Wild-type strain | [8] |
| *S. venezuelae* ATCC 10712 – pIJLK01 | *S. venezuelae* ATCC 10712 carrying the integrative plasmid pIJLK01, $Hyg^R$ | This study |
| *S. venezuelae* ATCC 10712 – pIJLK03 | *S. venezuelae* ATCC 10712 carrying the integrative plasmid pIJLK03, $Ksg^R$ | This study |
| *S. venezuelae* ATCC 10712 – pIJLK04 | *S. venezuelae* ATCC 10712 carrying the integrative plasmid pIJLK04, $Apr^R$ | This study |
| *S. venezuelae* ATCC 10712 – pIJLK05 | *S. venezuelae* ATCC 10712 carrying the integrative plasmid pIJLK05, $Sp^R$/$Sm^R$ | This study |
| *Streptomyces coelicolor* M145 | *S. coelicolor* A3(2) lacking plasmids SCP1 and SCP2 | [9] |
| *S. coelicolor* M145– pIJLK01 | *S. coelicolor* M145 carrying the integrative plasmid pIJLK01, $Hyg^R$ | This study |
| *S. coelicolor* M145– pIJLK03 | *S. coelicolor* M145 carrying the integrative plasmid pIJLK03, $Ksg^R$ | This study |
| *S. coelicolor* M145– pIJLK04 | *S. coelicolor* M145 carrying the integrative plasmid pIJLK04, $Apr^R$ | This study |
| *S. coelicolor* M145– pIJLK05 | *S. coelicolor* M145 carrying the integrative plasmid pIJLK05, $Sp^R$/$Sm^R$ | This study |
| *Streptomyces tenebrarius* ATCC 17920 | Wild-type strain | [10] |
| *Streptomyces kasugaensis* ATCC 15714 | Wild-type strain | [11] |

**Supplementary Table S3: Phages used in this study**

| Phage | Host organism | Lifestyle | Family | Genome | State of injected genome[12,13] | Reference |
|---|---|---|---|---|---|---|
| **Alderaan** | *S. venezuelae* ATCC 10712 | Virulent | *Siphoviridae* | dsDNA | Linear with terminal redundancy | [14] |
| **Coruscant** | *S. venezuelae* ATCC 10712 | Virulent | *Siphoviridae* | dsDNA | Linear with terminal repeats | [14] |
| **Dagobah** | *S. coelicolor* M145 | Temperate | *Siphoviridae* | dsDNA | Linear with terminal repeats | [14] |
| **Endor1** | *S. coelicolor* M145 | Temperate | *Siphoviridae* | dsDNA | Linear with terminal redundancy | [14] |
| **Endor2** | *S. coelicolor* M145 | Temperate | *Siphoviridae* | dsDNA | Linear with terminal redundancy | [14] |
| **CL31** | *C. glutamicum* MB001 | Temperate | *Siphoviridae* | dsDNA | Linear with cohesive ends | [15] |
| **Spe2** | *C. glutamicum* ATCC 13032 | Virulent | *Siphoviridae* | dsDNA | Unknown | This study, DSM110582 |
| **T4** | *E. coli* B (DSM613) | Virulent | *Myoviridae* | dsDNA | Linear with terminal redundancy | DSM4505 |
| **T5** | *E. coli* B (DSM613) | Virulent | *Siphoviridae* | dsDNA | Linear with terminal redundancy | DSM16353 |
| **T6** | *E. coli* B (DSM613) | Virulent | *Myoviridae* | dsDNA | Linear with terminal repeats | DSM4622 |
| **T7** | *E. coli* B (DSM613) | Virulent | *Podoviridae* | dsDNA | Linear with terminal repeats | DSM4623 |
| **M13** | *E. coli* W1485 (DSM5695) | Chronic infection | *Inoviridae* | ssDNA | Circular (+) strand | DSM13976 |
| **fd** | *E. coli* W1485 (DSM5695) | Chronic infection | *Inoviridae* | ssDNA | Circular (+) strand | DSM4498 |
| **MS2** | *E. coli* W1485 (DSM5695) | Virulent | *Leviviridae* | ssRNA | Linear, bound to the maturation protein [16] | DSM13767 |
| **Lambda (λ)** | *E. coli* LE392 (DSM4230) | Temperate | *Siphoviridae* | dsDNA | Linear with cohesive ends | DSM4499 |

**Supplementary Table S4: Plasmids used in this study**

| Plasmids | Characteristics | Reference |
|---|---|---|
| **pIJ10257** | Hyg$^R$; Cloning vector for the conjugal transfer of DNA from *E. coli* to *Streptomyces spp.;* (constitutive promoter *ermE**; Integration at the ΦBT1 attachment site) | [17] |
| **pEKEx2** | Kan$^R$; *C. glutamicum*/ *E. coli* shuttle vector for regulated gene expression; $P_{tac}$, *lacI*$^q$, pBL1 oriV$_{C.g.}$, pUC18 oriV$_{E.c.}$ | [18] |
| **pMKEx2** | Kan$^R$; *E. coli*/*C. glutamicum* shuttle vector based on pJC1 for expression of target genes under control of the $P_{T7}$ ($P_{lacI}$, *lacI*, $P_{T7}$, *lacO1*, N-term. Strep-tag II, MCS, C-term. His-tag, pHM1519 *ori*$_{C.g.}$; pACYC177 *ori*$_{E.c.}$) | [2] |
| **pMKEx-Kac** | Kan$^R$; Derivative of pEKEx2 with *kac338-C-strep* (kasugamycin resistance gene) under control of $P_{T7}$ | This study |
| **pIJ773** | pBluescript II SK(+)-based plasmid containing the apramycin resistance cassette flanked by FRT (FLP recognition target) recombination sites | [19] |
| **pCDFduet-1** | Sp$^R$/Sm$^R$; *E. coli* vector for coexpression of two target genes; $P_{T7}$, *lacI*, CloDF13 ori, T7 terminator | Novagen |
| **pUZ8002** | Kan$^R$; RP4 derivative with nontransmissible oriT | [20] |

| Plasmids | Characteristics | Template | Primer | Vector | Restriction enzyme | Sequencing primer | Reference |
|---|---|---|---|---|---|---|---|
| **pIJLK01** | Hyg$^R$; Derivative of pIJLK10257 with additional restrictions sites Bst1107I (upstream) and StuI (downstream) of the *aph(7")-Ia* gene allowing exchanging of the antibiotic cassette | pIJ10257 | 1 + 2<br>3 + 4<br>5 + 6 | pIJ10257 | KpnI;<br>PvuII | 31 - 34 | This study |
| **pIJLK03** | Ksg$^R$; Derivative of pIJLK01 with *aph(7")-Ia* exchanged for *kac338* (kasugamycin resistance gene) | pMKEx-Kac | 9 + 10 | pIJLK01 | Bst1107I;<br>StuI | 33 + 34 | This study |
| **pIJLK04** | Apr$^R$; Derivative of pIJLK01 with *aph(7")-Ia* exchanged for *aac(3)IV* (apramycin resistance gene) | pIJ773 | 11 + 12 | pIJLK01 | Bst1107I;<br>StuI | 33 + 34 | This study |

| | | | | | | | |
|---|---|---|---|---|---|---|---|
| **pIJLK05** | $Sp^R/Sm^R$; Derivative of pIJLK01 with *aph(7")-Ia* exchanged for *aadA* (spectinomycin/ streptomycin resistance gene) | pCDFduet-1 | 13 + 14 | pIJLK01 | Bst1107I; StuI | 33 + 34 | This study |
| **pEKEx2a** | $Kan^R$; Derivative of pEKEx2 with additional restrictions sites Bst1107I (upstream) and NotI (downstream) of the *aphA1* gene allowing exchanging of the antibiotic cassette | pEKEx2 | 15 + 16<br>17 + 18<br>19 + 20 | pEKEx2 | SapI; StuI | 35 – 37 | This study |
| **pEKEx2b** | $Hyg^R$; Derivative of pEKEx2a with *aphA1* exchanged for *aph(7")-Ia* (hygromycin resistance gene) | pIJ10257 | 21 + 22 | pEKEx2a | Bst1107I; NotI | 37 + 38 | This study |
| **pEKEx2c** | $Ksg^R$; Derivative of pEKEx2a with *aphA1* exchanged for *kac338* (kasugamycin resistance gene) | pMKEx-Kac | 23 + 24 | pEKEx2a | Bst1107I; NotI | 37 + 38 | This study |
| **pEKEx2d** | $Apr^R$; Derivative of pEKEx2a with *aphA1* exchanged for *aac(3)IV* (apramycin resistance gene) | pIJ773 | 25 + 26 | pEKEx2a | Bst1107I; NotI | 37 + 38 | This study |
| **pEKEx2e** | $Sp^R/Sm^R$; Derivative of pEKEx2a with *aphA1*exchanged for *aadA* (spectinomycin/ streptomycin resistance gene) | pCDFduet-1 | 27 + 28 | pEKEx2a | Bst1107I; NotI | 37 + 38 | This study |

| | | | | | | | |
|---|---|---|---|---|---|---|---|
| **pAN6_ aac(3)IV_Cstrep** | Kan$^{R}$; Derivative of pAN6 with *aac(3)IV* fused to a C-terminal Strep-tag | pIJ773 | 29+30 | pAN6_ CStrep | NdeI; NheI | 39 + 40 | This study |

**Supplementary Table S5: Oligonucleotides used in this study**

| No. | Oligonucleotide name | Sequence (5' - 3') |
|---|---|---|
| **Construction of plasmids** | | |
| **1** | pIJ10257_RE1_1_fw | TGCTCGGGTCGGGCTGGTACCAGTGAGCGTTTTTCAACCTCAG |
| **2** | pIJ10257_RE1_1_rv | GATTCTTGTGTCACGTATACAGCGGACCTCTATTCACAGGG |
| **3** | pIJ10257_RE1_2_fw | AATAGAGGTCCGCTGTATACGTGACACAAGAATCCCTGTTACTTCTCG |
| **4** | pIJ10257_RE2_2_rv | CGGGCGGCCCGGGGCGAGGCCTTCAGGCGCCGGGGG |
| **5** | pIJ10257_RE2_3_fw | CCCCCGGCGCCTGAAGGCCTCGCCCCGGGCCGC |
| **6** | pIJ10257_RE2_3_rv | GAAACCTGTCGTGCCAGCTGCATTAATGAATCGGCCAACGCGC |
| **7** | pIJLK02_kanR_fw | CCTGTGAATAGAGGTCCGCTGTATACATGAGCCATATTCAACGGGAAAC |
| **8** | pIJLK02_kanR_rv | GGCGGCCCGGGGCGAGGCCTTTAGAAAAACTCATCGAGCATCAAATGAA |
| **9** | pIJLK03_kac338_fw | AATAGAGGTCCGCTGTATACGTGCCGCGCTGGGCCG |
| **10** | pIJLK03_kac338_rv | GGCCCGGGGCGAGGCCTTTACAGGGCGATCAGCCCGG |
| **11** | pIJLK04_aac(3)IV_fw | TGAATAGAGGTCCGCTGTATACGTGCAATACGAATGGCGAAAAG |
| **12** | pIJLK04_aac(3)IV_rv | GGCGGCCCGGGGCGAGGCCTTCAGCCAATCGACTGGCG |
| **13** | pIJLK05_aadA_fw | TGAATAGAGGTCCGCTGTATACATGAGGGAAGCGGTGATCG |
| **14** | pIJLK05_aadA_rv | GCGGCCCGGGGCGAGGCCTTTATTTGCCGACTACCTTGGTGAT |
| **15** | pEKEx2_RE1_1_fw | GCGGTTTGCGTATTGGGCGCTCT |
| **16** | pEKEx2_RE1_1_rv | ATGGCTCATGTATACAACACCCCTTGTATTACTGTTTATGTAAGCAGAC |
| **17** | pEKEx2_RE1_2_fw | GGGGTGTTGTATACATGAGCCATATTCAACGGGAAACGTCT |
| **18** | pEKEx2_RE2_2_rv | TTCTGAGCGGCCGCTTAGAAAAACTCATCGAGCATCAAATGAAAC |
| **19** | pEKEx2_RE2_3_fw | TTTCTAAGCGGCCGCTCAGAATTGGTTAATTGGTTGTAACA |
| **20** | pEKEx2_RE2_3_rv | CGTGAAGAAGGTGTTGCTGACTC |
| **21** | pEKEx2b_hygR_fw | ATACAAGGGGTGTTGTATACGTGACACAAGAATCCCTGTTACTTCTC |
| **22** | pEKEx2b_hygR_rv | AATTAACCAATTCTGAGCGGCCGCTCAGGCGCCGGGGGC |
| **23** | pEKEx2c_kac338_fw | AATACAAGGGGTGTTGTATACGTGCCGCGCTGGGCCG |
| **24** | pEKEx2c_kac338_rv | ACCAATTCTGAGCGGCCGCTTACAGGGCGATCAGCCCG |
| **25** | pEKEx2d_aac(3)IV_fw | ATACAAGGGGTGTTGTATACGTGCAATACGAATGGCGAAAAG |
| **26** | pEKEx2d_aac(3)IV_rv | ACCAATTCTGAGCGGCCGCTCAGCCAATCGACTGGCGAG |
| **27** | pEKEx2e_aadA_fw | ACAAGGGGTGTTGTATACATGAGGGAAGCGGTGATCG |
| **28** | pEKEx2e_aadA_rv | ACCAATTCTGAGCGGCCGCTTATTTGCCGACTACCTTGGTGATC |
| **29** | pAN6_aac(3)IV_Cstrep_ fw | CCTGCAGAAGGAGATATACATATGATGTCATCAGCGGTGGAG |
| **30** | pAN6_aac(3)IV_CStrep_rv | TGTGGGTGGGACCAGCTAGCGCCAATCGACTGGCGAGC |
| **Sequencing primer** | | |
| **31** | pIJLK01_seq_fw | GATCAACCGCGACTAGCATC |
| **32** | pIJLK01_seq_rv | AGCAGTTCCGGGAAGAC |
| **33** | pIJLK0x_seq_fw | CGTAGAGATTGGCGATCCC |
| **34** | pIJLK0x_seq_rv | ATGCCAGGGCCTTTCAC |
| **35** | pEKEx2a_seq_fw | TTCCAGTCGGGAAACCTGTC |
| **36** | pEKEx2a_seq_rv | TCGCGAGCCCATTTATACCC |
| **37** | pEKEx2x_seq_rv | GCCTCGTGAAGAAGGTGTTG |
| **38** | pEKEx2x_seq_fw | GGAAAGCCACGTTGTGTCTC |
| **39** | pAN6_seq_ Cstrep_fw | CGGCGTTTCACTTCTGAGTTCGGC |
| **40** | pAN6_seq_ Cstrep_rv | GATATGACCATGATTACGCC |

| **qPCR primer** | | |
|---|---|---|
| **41** | qPCR_*atpD*_Sv_fw | TGTTCGAGACCGGCCTGAAG |
| **42** | qPCR_*atpD*_Sv_rv | AGACACCGTCGTGCAGCTTG |
| **43** | qPCR_Alderaan_fw | CTCGGCTATCCGATCATCC |
| **44** | qPCR_Alderaan_rv | TTGGTTGCGGTTGATGGAC |

**Supplementary Table S6: Polynucleotides used for phage targeting direct-geneFISH**

| No. | Sequence (5' - 3') |
|---|---|
| **Gene probes for phage targeting direct-geneFISH with Alderaan infecting *S. venezuelae*** | |
| **1** | ACGGCGATCAGCACCCAGGACAGGACGACGTCTTTGTGTCGCACCCACCATGGCGGTCTCGGTCTCGGCTCTTCCATCAACTTTCCCCCAGTTCGGCAACAGTCGACATCGTATGGAGAGAGGGGGTGAGCCCGTGCCCATTCACCCCGGCATGGTCGAGCCCCTCGCCGAACGCACCCGCGATCTCTACGCCGCCGCCG |
| **2** | TACGCGGACGCCGAGGACGCCTGTAACGGCTACCTCCTGAACAAGAAGGCCAAGGCGGACGGCATCAACCCGGCCGCCCTGTTCAGCGGCCCAGCCCGTATCGCGTACGCCCGAGCGTCGGACGAGCTGAAAGAGTGGTGGGCCGAACACGGTCGCCTAACGCAGGCGGAGTTCATCGAGCAGGTCACCGGCAAGGCTCA |
| **3** | CGCCGACAGCAACGGCCGTCACGAGCACGTCACCAACTACGACGTCGCCACGGCCTCCCCCACCACCCCGTAAGGAGGTCCGCCCATGGCGGGTAACAGCAGGTCCATCGACGCGCGCGGATGGCTCTTCGAGGTCAAGGACACCGACGCCAGCACCGAGACGTGGCTCCCGATCGCCGGTCTCAACTCCTGGTCGTACT |
| **4** | CGGCGCGTACGAGGAGGACGTCATGCAGCGCGGCGCCTCCATCACCCTGGAGGGTCAGTACCGCATCGACAAGACGACCAAGGCCCGCGACGTGGGACAGGCGTACATCGATGAGGAATGGACGCCGCGTCTCGGCATCGACTCGCACAACCAGATCCGCTACCGGCACGAGACGCAGTCCGCATGGGCGATCTGGGACG |
| **5** | CGTAGACGTCGCCGAAGACCTCTTGGACACGTTCGCCGAGTATCCGACGGAGTTCCGTCAGCTCGGCCTCGACGCCGAGACGGCGATGGGGCTGCTCTCCCAGGGACTCCAAGGCGGTGCGCGCGACGCCGATACGGTCGCCGACGCGTTGAAGGAATTCACGCTCATGGCTCAGGGCATGGGCGAGTCAACTGCGGAGT |
| **6** | GCCGTGTGGGCCGGGGCCATCGTCGTCGGTCAGTGGGTCCTCATGGCCACACAGGCCCTCATGCAGGCCGCCCGCATGGCCGCCGCGTGGCTCATCGCCATGGGCCCGATTGGCCTGCTCATCGCTGCCGTGGTCGGTCTCGTCGTCCTGATCATCAAGTATTGGGATGACATCGTCGCAGCCACCACGAAGGCGTGGGA |
| **7** | GGCCGCCGGCGACTACGTCGAGATGACCATTTTCAGCGGCGCCGCCCTGTCCGGCATTCCGTCCAGCTACAGCCGGGCGTCGTTGGTCTGGCAGGGCCCCGCGTGAGCGGCATGTACCGCGTCGTTCTGTGCGACCTGCGATCCGACCAAGTCCTCGACATCCTTCCCGCGCAGGGCATCAAGTGCGACGACTACATCGG |
| **8** | CCGCGACGTTCGACTCGTACCTCGCGCACCGGCTACTCAAGGACGGGTGGACCGGGAACGGGGTCGACCAACTCGACATCGCCCGTCAGATCGTCGACTGGGTCCAGTCGACCGAGGGCGGCAACATCGGCATCGAACTGGACTGGTCGCAGACATCCGGAGTGCTCCGCGACCGGGCGTACTCCCGCTACGACCTGTAC |
| **9** | GTCGTGCGCGACGTGCTCGACCAACTCGCCAACGTCGAGAACGGGTTCGAGTGGCGCGTACGTACGTACCGCGATGCGTCCGGCCGCCGCGTGAAGAGCTTGCAGCTCGGCTATCCGATCATCCGGAGCAGCCGTACCGAGTTGGTTCTCTCCTCCCCGGGCCCGGTCATCGACTACCGGATGCCCGAGGACGGCACCTC |
| **10** | GCGCCACCGCGCAGCTTCAGTTCCGCGTGAACGGAACCATCGTGGCGACCGGCACGGCGGGTCAACCGCTCCTTGCCACCTTCGCCATCCCGTCGTACGCGTTCGGCATGAACGCCGAGTTCGAGCTACAGGCCCGCGTGTCCAGCGGCACCGGAACCGCCTACGCCCAGACCCGCTACCTGTACGGCTTCCAGTCCTAA |
| **11** | GGCGCGGTGCTCGGCGCGGTGCTCGGCACGGTCGTCGTGGCGCTCATCGGCTGCTCTCCTCCTTGCGGGGCGTGCTCCGGAATGGCTCCACCTTGACACCCCGGTCCGGACGTATCTGCCCCCAAAGCGAGGACAGTTGAGACGTCTCTCGCTAGGGTCGAATAAGGCCCAAAAGTCCGGGCAGAGGGGGTTGACAGTGA |
| **12** | AACCGCGCGGACGCTGCACGGGCGCTGGGAGTGGATGAGGAAATGATCTGGCCGAAGGCGGTGCAGGACCGCGTAAAGGTCGGCGGCGACCGGGAGATCCTCCGCACTTACCCATACCGCTCGGCGTGCCCCTCCAACGTGTGGGCGGACCTCGCTGCCGGCGCCGAGCACGAGCTGTTTCTCGCCGGGTATACGAACTA |
| **13** | GCGAGGTGACCCGGCAGCGCGAGGTAATCGAAGGCGTTCCGCTGTCGGTTTCCACGCGCATTCGGATCACGCTCGATGAGTTGGCGCGGCTCGGGTCGGTCGAGGGTGTCGAGGCCCGGCTGAGCGCTGCCGAGGATGCCGTAAATCACGTGAGCCTGTCGGTATTCCGATTCGACGAGGAGGCCCTCGTAACGCCTCAT |
| **14** | TGCGGCGGCACGTTGACGGCGGGATGTTCGACCGCTTCGCAGAGCACGCCGAAGAGCTGTGGGAGCGGGCCGTGCCCGTGACGTCGTAGACGACGAACGGCCCCGCCTCCCAGACAGGGACGCGGGGCCGTCGCGTTTCCGCAGGTCAAGCAACCATTCCGTATCGCTCCCCTTTAGGTCCACTGTCGTGGGGTGTTGGA |

| | |
|---|---|
| **15** | GACGCGAGTGTCTGTGACTGCTTCGCCCCTGAGGAGCCGGCCGACGGTCGAGGCGCTGATGCCGCTATCTGCGGCAAAGCGGGAGCGCCCCCCGCTGCGTAGGCCGGTGACGTCGTATCCGCGTCTCACGAGTTCCTGGGAGAGCCATGCGGCGAACGCCGAGCGCGTCGCGTCGGTCTTGTTTTCTGCCATGGCAGAAA |
| **16** | CCGCGCGGCGAGAAGGGCACGCACACCGTGCTCCGGATGCTGTACCGCATCTACGGCCCGGCCGGCGACTGCCTCGCCGTCACCATCCCGGGCGAGGCCATGGACACGGCCGACAAGAGCACCAACAAGGCGATGTCGGCCGCGCTCAAGTACATGCTCTTTCAGGTGTTCATGATCCCCGTGGACGCCCGCAGCATCGA |
| **17** | GGTGATCCCGCCCGCGTTCACCCCGGAGGACGGCACCTAGCCCCCCTCGATAGGGGGAGCCGGATCAGCCCCACATCCGCTATCTTTCTGTCTCGACAGATACATGCCTGTCGAGACAGAACCGCGAGCGCGCGCGCTCAGCACGTAGAGCAGGAGGACCGCCCCGTGAACACCCCCGAGCGCTTCGCCGCCAAGGTCGA |
| **Gene probes for phage targeting direct-geneFISH with λ infecting *E. coli*** | |
| **1** | AGCAGTATCTTAAATTTGGCGACAAAGAGACGCCGTTTGGCCTCAAATGGACGCCGGATGACCCCTCCAGCGTGTTTTATCTCTGCGAGCATAATGCCTGCGTCATCCGCCAGCAGGAGCTGGACTTTACTGATGCCCGTTATATCTGCGAAAAGACCGGGATCTGGACCCGTGATGGCATTCTCTGGTTTTCGTCATCCGGTGAAGAGATTGAGCCACCTGACAGTGTGACCTTTCACATCTGGACAGCGTACAGCCCGTTCACCACCTGGGTGCAGATTGTCAAAGACTGGATGAAAA |
| **2** | GCAGGACAACGTATTCGATGTGTTATCTGAAAGTACTGATGAACGGTGCGGTGATTTATGATGGCGCGGCGAACGAGGCGGTACAGGTGTTCTCCCGTATTGTTGACATGCCAGCGGGTCGGGGAAACGTGATCCTGACGTTCACGCTTACGTCCACACGGCATTCGGCAGATATTCCGCCGTATACGTTTGCCAGCGATGTGCAGGTTATGGTGATTAAGAAACAGGCGCTGGGCATCAGCGTGGTCTGAGTGTGTTACAGAGGTTCGTCCGGGAACGGGCGTTTTATTATAAAACAGT |
| **3** | AGATTATTATGGGCCGCCACGACGATGAACAGACGCTGCTGCGTGTGGATGAGGCCATCAATAAAACCTATACCCGCCGGAATGGTGCAGAAATGTCGATATCCCGTATCTGCTGGGATACTGGCGGGATTGACCCGACCATTGTGTATGAACGCTCGAAAAAACATGGGCTGTTCCGGGTGATCCCCATTAAAGGGGCATCCGTCTACGGAAAGCCGGTGGCCAGCATGCCACGTAAGCGAAACAAAAACGGGGTTTACCTTACCGAAATCGGTACGGATACCGCGAAAGAGCAGATTT |
| **4** | CAATTTTGTCCCACTCCCTGCCTCTGTCATCACGATACTGTGATGCCATGGTGTCCGACTTATGCCCGAGAAGATGTTGAGCAAACTTATCGCTTATCTGCTTCTCATAGAGTCTTGCAGACAAACTGCGCAACTCGTGAAAGGTAGGCGGATCCCCTTCGAAGGAAAGACCTGATGCTTTTCGTGCGCGCATAAAATACCTTGATACTGTGCCGGATGAAAGCGGTTCGCGACGAGTAGATGCAATTATGGTTTCTCCGCCAAGAATCTCTTTGCATTTATCAAGTGTTTCCTTCATTG |
| **5** | TGCTCGACATAAAGATATCCATCTACGATATCAGACCACTTCATTTCGCATAAATCACCAACTCGTTGCCCGGTAACAACAGCCAGTTCCATTGCAAGTCTGAGCCAACATGGTGATGATTCTGCTGCTTGATAAATTTTCAGGTATTCGTCAGCCGTAAGTCTTGATCTCCTTACCTCTGATTTTGCTGCGCGAGTGGCAGCGACATGGTTTGTTGTTATATGGCCTTCAGCTATTGCCTCTCGGAATGCATCGCTCAGTGTTGATCTGATTAACTTGGCTGACGCCGCCTTGCCCTCG |
| **6** | AACTCAATGTTGGCCTGTATAGCTTCAGTGATTGCGATTCGCCTGTCTCTGCCTAATCCAAACTCTTTACCCGTCCTTGGGTCCCTGTAGCAGTAATATCCATTGTTTCTTATATAAAGGTTAGGGGGTAAATCCCGGCGCTCATGACTTCGCCTTCTTCCCATTTCTGATCCTCTTCAAAAGGCCACCTGTTACTGGTCGATTTAAGTCAACCTTTACCGCTGATTCGTGGAACAGATACTCTCTTCCATCCTTAACCGGAGGTGGGAATATCCTGCATTCCCGAACCCATCGACGAAC |
| **7** | TGTTTCAAGGCTTCTTGGACGTCGCTGGCGTGCGTTCCACTCCTGAAGTGTCAAGTACATCGCAAAGTCTCCGCAATTACACGCAAGAAAAAACCGCCATCAGGCGGCTTGGTGTTCTTTCAGTTCTTCAATTCGAATATTGGTTACGTCTGCATGTGCTATCTGCGCCCATATCATCCAGTGGTCGTAGCAGTCGTTGATGTTCTCCGCTTCGATAACTCTGTTGAATGGCTCTCCATTCCATTCTCCTGTGACTCGGAAGTGCATTTATCATCTCCATAAAACAAAACCCGCCGTAGC |
| **8** | ACTCAACCCGATGTTTGAGTACGGTCATCATCTGACACTACAGACTCTGGCATCGCTGTGAAGACGACGCGAAATTCAGCATTTTCACAAGCGTTATCTTTTACAAAACCGATCTCACTCTCCTTTGATGCGAATGCCAGCGTCAGACATCATATGCAGATACTCACCTGCATCCTGAACCCATTGACCTCCAACCCCGTAATAGCGATGCGTAATGATGTCGATAGTTACTAACGGGTCTTGTTCGATTAACTGCCGCAGAAACTCTTCCAGGTCACCAGTGCAGTGCTTGATAACAGG |
| **9** | GTTCATCCAGCAGTTCCAGCACAATCGATGGTGTTACCAATTCATGGAAAAGGTCTGCGTCAAATCCCCAGTCGTCATGCATTGCCTGCTCTGCCGCTTCACGCAGTGCCTGAGAGTTAATTTCGCTCACTTCGAACCTCTCTGTTTACTGATAAGTTCCAGATCCTCCTGGCAACTTGCACAAGTCCGACAACCCTGAACGACCAGGCGTCTTCGTTCATCTATCGGATCGCCACACTCACAACAATGAGTGGCAGATATAGCCTGGTGGTTCAGGCGGCGCATTTTTATTGCTGTGTT |

| | |
|---|---|
| **10** | TGAGGGTGAATGCGAATAATAAAAAAGGAGCCTGTAGCTCCCTGATGATTTTGCTTTTCATGTT CATCGTTCCTTAAAGACGCCGTTTAACATGCCGATTGCCAGGCTTAAATGAGTCGGTGTGAAT CCCATCAGCGTTACCGTTTCGCGGTGCTTCTTCAGTACGCTACGGCAAATGTCATCGACGTTT TTATCCGGAAACTGCTGTCTGGCTTTTTTTGATTTCAGAATTAGCCTGACGGGCAATGCTGCGA AGGGCGTTTTCCTGCTGAGGTGTCATTGAACAAGTCCCATGTCGGC |
| **11** | AGGTAAACGGGCATTTCAGTTCAAGGCCGTTGCCGTCACTGCATAAACCATCGGGAGAGCAG GCGGTACGCATACTTTCGTCGCGATAGATGATCGGGGATTCAGTAACATTCACGCCGGAAGTG AATTCAAACAGGGTTCTGGCGTCGTTCTCGTACTGTTTTCCCCAGGCCAGTGCTTTAGCGTTAA CTTCCGGAGCCACACCGGTGCAAACCTCAGCAAGCAGGGTGTGGAAGTAGGACATTTTCATG TCAGGCCACTTCTTTCCGGAGCGGGGTTTTGCTATCACGTTGTGAACTT |
| **12** | TGATGACGCCGAGCCGTAATTTGTGCCACGCATCATCCCCCTGTTCGACAGCTCTCACATCGA TCCCGGTACGCTGCAGGATAATGTCCGGTGTCATGCTGCCACCTTCTGCTCTGCGGCTTTCTG TTTCAGGAATCCAAGAGCTTTTACTGCTTCGGCCTGTGTCAGTTCTGACGATGCACGAATGTC GCGGCGAAATATCTGGGAACAGAGCGGCAATAAGTCGTCATCCCATGTTTTATCCAGGGCGAT CAGCAGAGTGTTAATCTCCTGCATGGTTTCATCGTTAACCGGAGTGAT |
| **13** | TCGCGTTCCGGCTGACGTTCTGCAGTGTATGCAGTATTTTCGACAATGCGCTCGGCTTCATCC TTGTCATAGATACCAGCAAATCCGAAGGCCAGACGGGCACACTGAATCATGGCTTTATGACGT AACATCCGTTTGGGATGCGACTGCCACGGCCCCGTGATTTCTCTGCCTTCGCGAGTTTTGAAT GGTTCGCGGCGGCATTCATCCATCCATTCGGTAACGCAGATCGGATGATTACGGTCCTTGCG GTAAATCCGGCATGTACAGGATTCATTGTCCTGCTCAAAGTCCATGCCA |
| **14** | TCAAACTGCTGGTTTTCATTGATGATGCGGGACCAGCCATCAACGCCCACCACCGGAACGATG CCATTCTGCTTATCAGGAAAGGCGTAAATTTCTTTCGTCCACGGATTAAGGCCGTACTGGTTG GCAACGATCAGTAATGCGATGAACTGCGCATCGCTGGCATCACCTTTAAATGCCGTCTGGCGA AGAGTGGTGATCAGTTCCTGTGGGTCGACAGAATCCATGCCGACACGTTCAGCCAGCTTCCC AGCCAGCGTTGCGAGTGCAGTACTCATTCGTTTTATACCTCTGAATCAA |
| **15** | TATCAACCTGGTGGTGAGCAATGGTTTCAACCATGTACCGGATGTGTTCTGCCATGCGCTCCT GAAACTCAACATCGTCATCAAACGCACGGGTAATGGATTTTTTGCTGGCCCCGTGGCGTTGCA AATGATCGATGCATAGCGATTCAAACAGGTGCTGGGGCAGGCCTTTTTCCATGTCGTCTGCCA GTTCTGCCTCTTTCTCTTCACGGGCGAGCTGCTGGTAGTGACGCGCCCAGCTCTGAGCCTCA AGACGATCCTGAATGTAATAAGCGTTCATGGCTGAACTCCTGAAATAGC |
| **16** | GATAAAGCCAAGGCCAATATCTAAGTAACTAGATAAGAGGAATCGATTTTCCCTTAATTTTCTG GCGTCCACTGCATGTTATGCCGCGTTCGCCAGGCTTGCTGTACCATGTGCGCTGATTCTTGCG CTCAATACGTTGCAGGTTGCTTTCAATCTGTTTGTGGTATTCAGCCAGCACTGTAAGGTCTATC GGATTTAGTGCGCTTTCTACTCGTGATTTCGGTTTGCGATTCAGCGAGAGAATAGGGCGGTTA ACTGGTTTTGCGCTTACCCCAACCAACAGGGGATTTGCTGCTTTCC |
| **17** | AGCCTGTTTCTCTGCGCGACGTTCGCGGCGGCGTGTTTGTGCATCCATCTGGATTCTCCTGTC AGTTAGCTTTGGTGGTGTGTGGCAGTTGTAGTCCTGAACGAAAACCCCCCGCGATTGGCACAT TGGCAGCTAATCCGGAATCGCACTTACGGCCAATGCTTCGTTTCGTATCACACACCCCAAAGC CTTCTGCTTTGAATGCTGCCCTTCTTCAGGGCTTAATTTTTAAGAGCGTCACCTTCATGGTGGT CAGTGCGTCCTGCTGATGTGCTCAGTATCACCGCCAGTGGTATTTAT |
| **18** | TACTATGTTATGTTCTGAGGGGAGTGAAAATTCCCCTAATTCGATGAAGATTCTTGCTCAATTG TTATCAGCTATGCGCCGACCAGAACACCTTGCCGATCAGCCAAACGTCTCTTCAGGCCACTGA CTAGCGATAACTTTCCCCACAACGGAACAACTCTCATTGCATGGGATCATTGGGTACTGTGGG TTTAGTGGTTGTAAAAACACCTGACCGCTATCCCTGATCAGTTTCTTGAAGGTAAACTCATCAC CCCCAAGTCTGGCTATGCAGAAATCACCTGGCTCAACAGCCTGCTC |
| **19** | TATTTGCATACATTCAATCAATTGTTATCTAAGGAAATACTTACATATGGTTCGTGCAAACAAAC GCAACGAGGCTCTACGAATCGAGAGTGCGTTGCTTAACAAAATCGCAATGCTTGGAACTGAGA AGACAGCGGAAGCTGTGGGCGTTGATAAGTCGCAGATCAGCAGGTGGAAGAGGGACTGGATT CCAAAGTTCTCAATGCTGCTTGCTGTTCTTGAATGGGGGGTCGTTGACGACGACATGGCTCGA TTGGCGCGACAAGTTGCTGCGATTCTCACCAATAAAAAACGCCCGGC |
| **20** | TCAAGCAGCAAGGCGGCATGTTTGGACCAAATAAAAACATCTCAGAATGGTGCATCCCTCAAA ACGAGGGAAAATCCCCTAAAACGAGGGATAAAACATCCCTCAAATTGGGGGATTGCTATCCCT CAAAACAGGGGGACACAAAAGACACTATTACAAAAGAAAAAAGAAAAGATTATTCGTCAGAGAA TTCTGGCGAATCCTCTGACCAGCCAGAAAACGACCTTTCTGTGGTGAAACCGGATGCTGCAAT TCAGAGCGGCAGCAAGTGGGGGACAGCAGAAGACCTGACCGCCGCAG |
| **21** | ATAAGTGGACCCAACTCGAAATCAACCGTAACAAGCAACAGGCAGGCGTGACAGCCAGCAAA CCAAAACTCGACCTGACAAACACAGACTGGATTTACGGGGTGGATCTATGAAAAACATCGCCG CACAGATGGTTAACTTTGACCGTGAGCAGATGCGTCGGATCGCCAACAACATGCCGGAACAG TACGACGAAAAGCCGCAGGTACAGCAGGTAGCGCAGATCATCAACGGTGTGTTCAGCCAGTT ACTGGCAACTTTCCCGGCGAGCCTGGCTAACCGTGACCAGAACGAAGTGAA |

| | |
|---|---|
| **22** | TGGCGGTATATGGAGTTAAAAGATGACCATCTACATTACTGAGCTAATAACAGGCCTGCTGGT AATCGCAGGCCTTTTTATTTGGGGGAGAGGGAAGTCATGAAAAAACTAACCTTTGAAATTCGAT CTCCAGCACATCAGCAAAACGCTATTCACGCAGTACAGCAAATCCTTCCAGACCCAACCAAAC CAATCGTAGTAACCATTCAGGAACGCAACCGCAGCTTAGACCAAAACAGGAAGCTATGGGCCT GCTTAGGTGACGTCTCTCGTCAGGTTGAATGGCATGGTCGCTGGCTG |
| **23** | GATGCAGAAAGCTGGAAGTGTGTGTTTACCGCAGCATTAAAGCAGCAGGATGTTGTTCCTAAC CTTGCCGGGAATGGCTTTGTGGTAATAGGCCAGTCAACCAGCAGGATGCGTGTAGGCGAATT TGCGGAGCTATTAGAGCTTATACAGGCATTCGGTACAGAGCGTGGCGTTAAGTGGTCAGACG AAGCGAGACTGGCTCTGGAGTGGAAAGCGAGATGGGGAGACAGGGCTGCATGATAAATGTCG TTAGTTTCTCCGGTGGCAGGACGTCAGCATATTTGCTCTGGCTAATGGAGC |
| **24** | CCATTTCGGGCGAGGGAATTACACCACGTGGATTGGCATCAGAGCTGATGAACCGAAGCGGC TAAAGCCAAAGCCTGGAATCAGATATCTTGCTGAACTGTCAGACTTTGAGAAGGAAGATATCCT CGCATGGTGGAAGCAACAACCATTCGATTTGCAAATACCGGAACATCTCGGTAACTGCATATT CTGCATTAAAAAATCAACGCAAAAAATCGGACTTGCCTGCAAAGATGAGGAGGGATTGCAGCG TGTTTTTAATGAGGTCATCACGGGATCCCATGTGCGTGACGGACATCG |
| **25** | GGAAACGCCAAAGGAGATTATGTACCGAGGAAGAATGTCGCTGGACGGTATCGCGAAAATGT ATTCAGAAAATGATTATCAAGCCCTGTATCAGGACATGGTACGAGCTAAAAGATTCGATACCGG CTCTTGTTCTGAGTCATGCGAAATATTTGGAGGGCAGCTTGATTTCGACTTCGGGAGGGAAGC TGCATGATGCGATGTTATCGGTGCGGTGAATGCAAAGAAGATAACCGCTTCCGACCAAATCAA CCTTACTGGAATCGATGGTGTCTCCGGTGTGAAAGAACACCAACAGGG |
| **26** | GTGTTACCACTACCGCAGGAAAAGGAGGACGTGTGGCGAGACAGCGACGAAGTATCACCGAC ATAATCTGCGAAAACTGCAAATACCTTCCAACGAAACGCACCAGAAATAAACCCAAGCCAATCC CAAAAGAATCTGACGTAAAAACCTTCAACTACACGGCTCACCTGTGGGATATCCGGTGGCTAA GACGTCGTGCGAGGAAAACAAGGTGATTGACCAAAATCGAAGTTACGAACAAGAAAGCGTCG AGCGAGCTTTAACGTGCGCTAACTGCGGTCAGAAGCTGCATGTGCTGGA |
| **27** | GCGCAGAACTGATGAGCGATCCGAATAGCTCGATGCACGAGGAAGAAGATGATGGCTAAACC AGCGCGAAGACGATGTAAAAACGATGAATGCCGGGAATGGTTTCACCCTGCATTCGCTAATCA GTGGTGGTGCTCTCCAGAGTGTGGAACCAAGATAGCACTCGAACGACGAAGTAAAGAACGCG AAAAAGCGGAAAAAGCAGCAGAGAAGAAACGACGACGAGAGGAGCAGAAACAGAAAGATAAA CTTAAGATTCGAAAACTCGCCTTAAAGCCCCGCAGTTACTGGATTAAACAA |
| **28** | CCAACAAGCCGTAAACGCCTTCATCAGAGAAAGAGACCGCGACTTACCATGTATCTCGTGCGG AACGCTCACGTCTGCTCAGTGGGATGCCGGACATTACCGGACAACTGCTGCGGCACCTCAAC TCCGATTTAATGAACGCAATATTCACAAGCAATGCGTGGTGTGCAACCAGCACAAAAGCGGAA ATCTCGTTCCGTATCGCGTCGAACTGATTAGCCGCATCGGGCAGGAAGCAGTAGACGAAATC GAATCAAACCATAACCGCCATCGCTGGACTATCGAAGAGTGCAAGGCGAT |
| **29** | TGTTATCTGCCACGCCGATTATCCCTTTGACGAATACGAGTTTGGAAAGCCAGTTGATCATCAG CAGGTAATCTGGAACCGCGAACGAATCAGCAACTCACAAAACGGGATCGTGAAAGAAATCAAA GGCGCGGACACGTTCATCTTTGGTCATACGCCAGCAGTGAAACCACTCAAGTTTGCCAACCAA ATGTATATCGATACCGGCGCAGTGTTCTGCGGAAACCTAACATTGATTCAGGTACAGGGAGAA GGCGCATGAGACTCGAAAGCGTAGCTAAATTTCATTCGCCAAAAAGC |
| **30** | CAGAGATTGCCATGGTACAGGCCGTGCGGTTGATATTGCCAAAACAGAGCTGTGGGGGAGAG TTGTCGAGAAAGAGTGCGGAAGATGCAAAGGCGTCGGCTATTCAAGGATGCCAGCAAGCGCA GCATATCGCGCTGTGACGATGCTAATCCCAAACCTTACCCAACCCACCTGGTCACGCACTGTT AAGCCGCTGTATGACGCTCTGGTGGTGCAATGCCACAAAGAAGAGTCAATCGCAGACAACATT TTGAATGCGGTCACACGTTAGCAGCATGATTGCCACGGATGGCAACATAT |
| **31** | TGAATAAAATTGGGTAAATTTGACTCAACGATGGGTTAATTCGCTCGTTGTGGTAGTGAGATGA AAAGAGGCGGCGCTTACTACCGATTCCGCCTAGTTGGTCACTTCGACGTATCGTCTGGAACTC CAACCATCGCAGGCAGAGAGGTCTGCAAAATGCAATCCCGAAACAGTTCGCAGGTAATAGTTA GAGCCTGCATAACGGTTTCGGGATTTTTTATATCTGCACAACAGGTAAGAGCATTGAGTCGATA ATCGTGAAGAGTCGGCGAGCCTGGTTAGCCAGTGCTCTTTCCGTTG |
| **32** | TGCTGAATTAAGCGAATACCGGAAGCAGAACCGGATCACCAAATGCGTACAGGCGTCATCGC CGCCCAGCAACAGCACAACCCAAACTGAGCCGTAGCCACTGTCTGTCCTGAATTCATTAGTAA TAGTTACGCTGCGGCCTTTTACACATGACCTTCGTGAAAGCGGGTGGCAGGAGGTCGCGCTA ACAACCTCCTGCCGTTTTGCCCGTGCATATCGGTCACGAACAAATCTGATTACTAAACACAGTA GCCTGGATTTGTTCTATCAGTAATCGACCTTATTCCTAATTAAATAGAG |
| **33** | CAAATCCCCTTATTGGGGGTAAGACATGAAGATGCCAGAAAAACATGACCTGTTGGCCGCCAT TCTCGCGGCAAAGGAACAAGGCATCGGGGCAATCCTTGCGTTTGCAATGGCGTACCTTCGCG GCAGATATAATGGCGGTGCGTTTACAAAAACAGTAATCGACGCAACGATGTGCGCCATTATCG CCTGGTTCATTCGTGACCTTCTCGACTTCGCCGGACTAAGTAGCAATCTCGCTTATATAACGAG CGTGTTTATCGGCTACATCGGTACTGACTCGATTGGTTCGCTTATCAA |

| | |
|---|---|
| **34** | ATCATGGTTATGACGTCATTGTAGGCGGAGAGCTATTTACTGATTACTCCGATCACCCTCGCAA<br>ACTTGTCACGCTAAACCCAAAACTCAAATCAACAGGCGCCGGACGCTACCAGCTTCTTTCCCG<br>TTGGTGGGATGCCTACCGCAAGCAGCTTGGCCTGAAAGACTTCTCTCCGAAAAGTCAGGACG<br>CTGTGGCATTGCAGCAGATTAAGGAGCGTGGCGCTTTACCTATGATTGATCGTGGTGATATCC<br>GTCAGGCAATCGACCGTTGCAGCAATATCTGGGCTTCACTGCCGGGCG |
| **35** | GATAAAACAAAAGCCACCGTGTCGGTCAGTGGTATGACCATCACCGTGAACGGCGTTGCTGC<br>AGGCAAGGTCAACATTCCGGTTGTATCCGGTAATGGTGAGTTTGCTGCGGTTGCAGAAATTAC<br>CGTCACCGCCAGTTAATCCGGAGAGTCAGCGATGTTCCTGAAAACCGAATCATTTGAACATAA<br>CGGTGTGACCGTCACGCTTTCTGAACTGTCAGCCCTGCAGCGCATTGAGCATCTCGCCCTGAT<br>GAAACGGCAGGCAGAACAGGCGGAGTCAGACAGCAACCGGAAGTTTACT |

### Figures

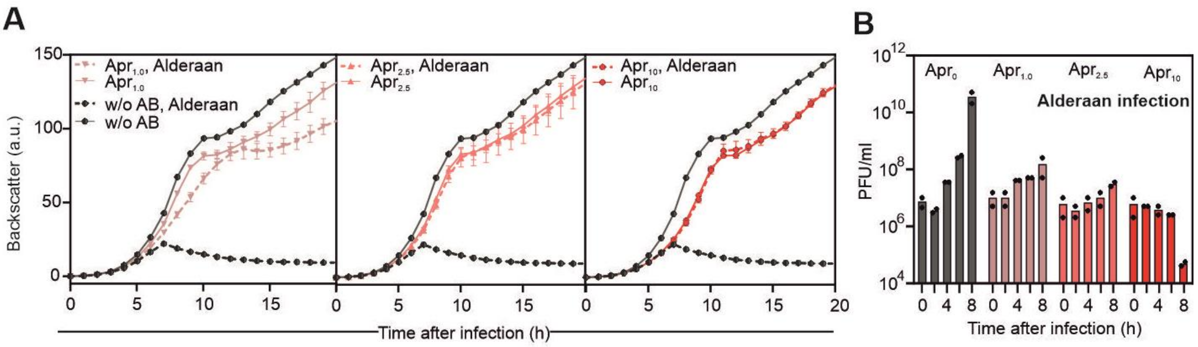


**Supplementary Figure S1 | Dose-dependent effect of apramycin on the *Streptomyces* phage Alderaan. A**, Growth of *Streptomyces venezuelae* infected with the phage Alderaan showing the dose-dependent effects of apramycin on infection (n = 3 independent biological replicates; error bars represent s.d.). **B,** Corresponding phage titers over time in presence of increasing concentrations of apramycin (0, 1, 2.5 and 10 µg/ml). Data represent an average of two independent biological replicates.

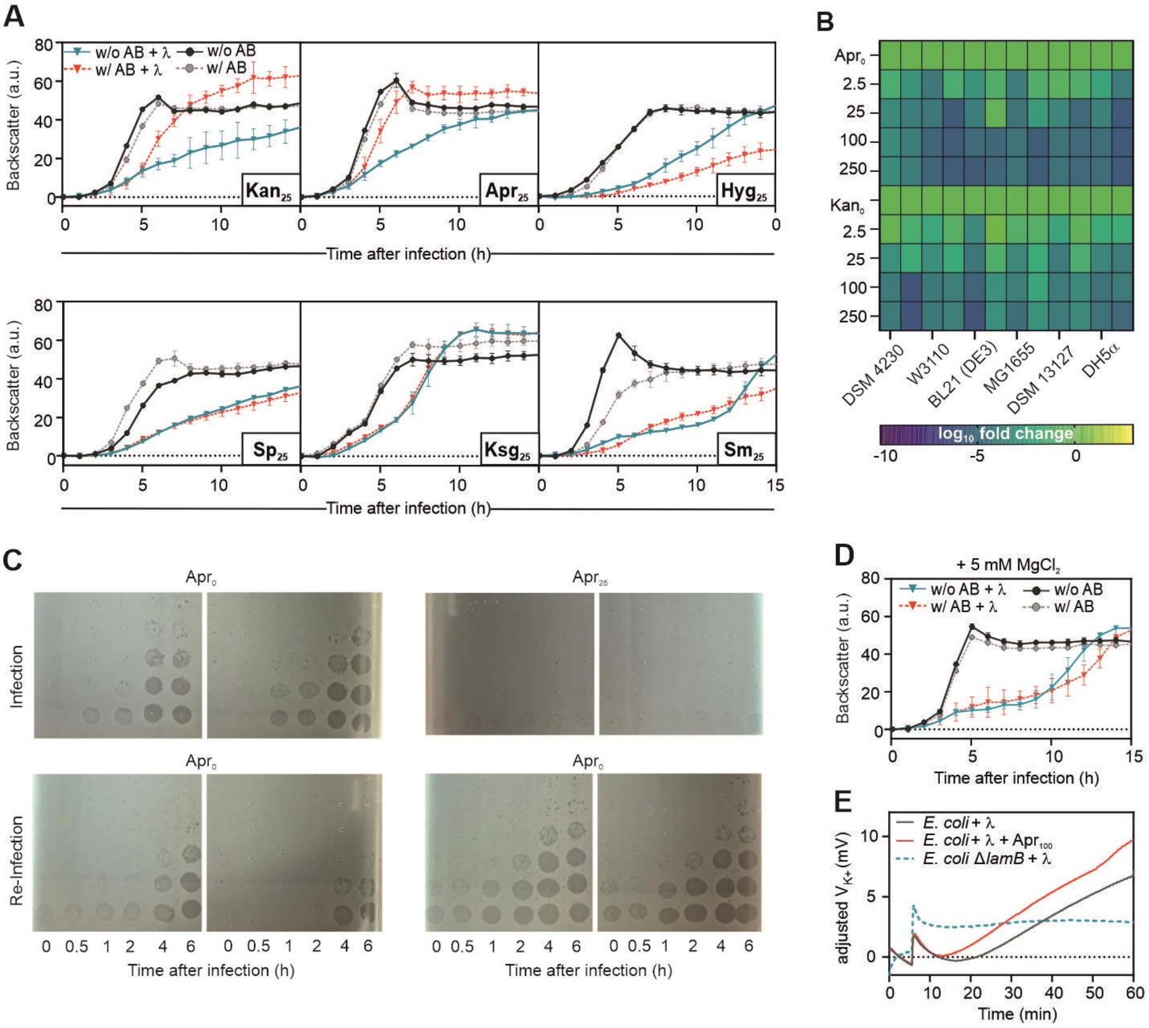


**Supplementary Figure S2 | Effect of aminoglycosides on *E. coli* phage λ. A**, Infection curves of *E. coli* infected with phage λ in presence of different aminoglycosides (n = 3 independent biological replicates; error bars represent s. d.). **B,** Heat map showing the $log_{10}$-fold change in plaque formation by λ on different *E. coli* strains in the presence of aminoglycosides relative to the aminoglycoside-free control. **C**, Re-infection of cultures previously treated with apramycin ($Apr_{25}$, upper row), shows efficient infection of *E. coli* by phage λ in the absence of apramycin ($Apr_{25}$, lower row)**. D,** Addition of $MgCl_2$ counteracts the effect of apramycin on infection of *E. coli* by λ. **E**, Potassium efflux assays performed with *E. coli* WT and the Δ*lamB* strain (lacking the λ receptor). λ was added after 5.5 minutes.

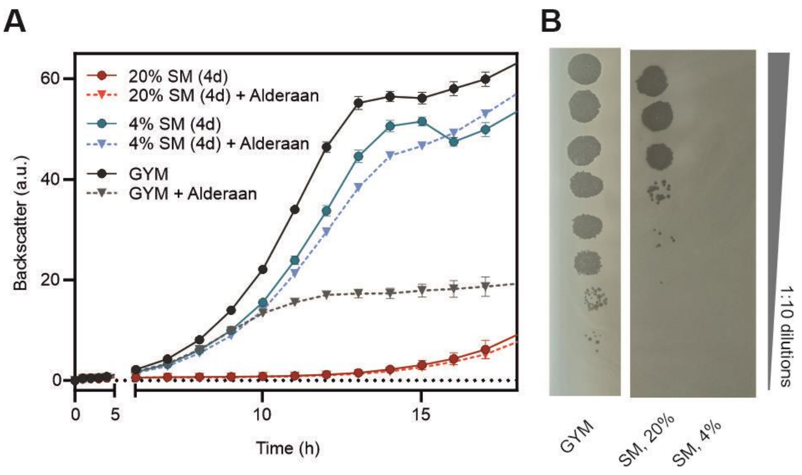


**Supplementary Figure S3 | Secondary metabolites produced by *S. kasugaensis* inhibit phage infection. A**, Influence of spent medium (SM) from *S. kasugaensis* (harvested after 4 days of cultivation) on infection of *S. venezuelae* by Alderaan (n = 3 independent biological replicates; error bars represent s. d.). **B**, Determination of the final phage titers of infected cultures shown in **A.** Results are representative of two replicates.

Please note, that the PFU observed for the sample with 20% represents the initial phage titer added to the culture, which is retained due to the inhibition of bacterial growth. In contrast, the phage titer declined to zero in the cultures where 4% of conditioned medium was added, since infection of growing cells occurred but phages were not able to complete their life cycle.

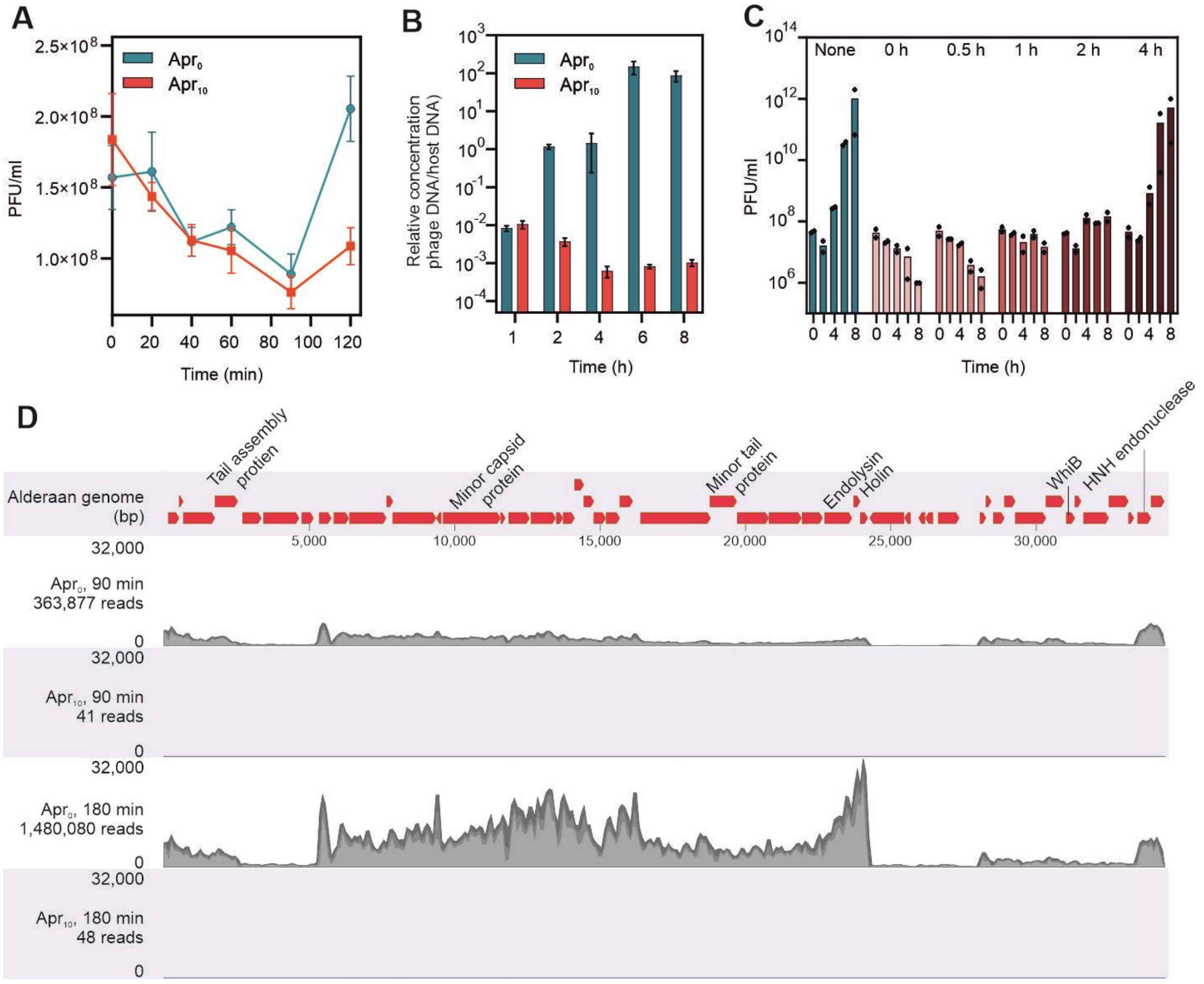


**Supplementary Figure S4 | Investigations of the mechanism of action of apramycin. A**, Effect of apramycin on phage adsorption of phage Alderaan to *S. venezuelae*. Shown is the time-resolved quantification of extracellular Alderaan DNA via qPCR. Culture supernatants were pre-treated with 100 U/ml DNase to exclusively quantify phage DNA deriving from intact phage particles. A DNase-treated phage stock with known phage titer was used to infer phage titers (in PFU/ml) from DNA quantification. **B**, Time-resolved quantification of intracellular Alderaan DNA via qPCR. Data of **A** and **B** represent mean values of two independent biological replicates measured as technical duplicates. **C**, Impact of apramycin when added at different time points post phage infection. For each sample, phage titers were measured over time. Data represent an average of 2 independent biological replicates. **D**, Enlargment of **Fig.4D** showing the RNA-seq coverage of the Alderaan genome in presence or absence of apramycin. Genome organization of Alderaan is displayed in the upper part.

**Videos**

**Video S1: Apramycin prevents cell lysis during infection of *S. venezuelae* with phage Alderaan.** Time-lapse video of *S. venezuelae* ATCC 10712 carrying the pIJLK04 plasmid, which was cultivated in a microfluidics system and challenged with Alderaan ($10^8$ PFU/ml; flow rate 200 nl/min) in presence and absence of 5 or 10 µg/ml apramycin.

**The video is provided as a separate file:**

- Video S1_Svenezueale_Alderaan_apramycin effect
